# APOBEC3A Catalyzes Mutation and Drives Carcinogenesis *In Vivo*

**DOI:** 10.1101/2019.12.27.889345

**Authors:** Emily K. Law, Rena Levin-Klein, Matthew C. Jarvis, Hyoung Kim, Prokopios P. Argyris, Michael A. Carpenter, Gabriel J. Starrett, Lindsay K. Larson, Michael B. Burns, Rachel I. Vogel, Spyridon Stavrou, Alexya N. Aguilera, Sandra Wagner, David A. Largaespada, Timothy K. Starr, Susan R. Ross, Reuben S. Harris

## Abstract

The APOBEC3 family of antiviral DNA cytosine deaminases is implicated as the second largest source of mutation in cancer. This mutational process may be a causal driver or inconsequential passenger to the overall tumor phenotype. We show that human APOBEC3A expression in murine colon and liver tissues increases tumorigenesis. All other APOBEC3 family members including APOBEC3B did not as strongly promote liver tumor formation. DNA sequences from APOBEC3A-expressing animals display hallmark APOBEC signature mutations in TCA/T motifs. Bioinformatic comparisons of the observed APOBEC3A mutation signature in murine tumors, previously reported APOBEC3A and APOBEC3B mutation signatures in yeast, and reanalyzed APOBEC mutation signatures in human tumor data sets support cause-and-effect relationships for APOBEC3A-catalyzed deamination and mutagenesis in driving multiple human cancers.

## INTRODUCTION

DNA damage and mutation are fundamental hallmarks of cancer (Hanahan and Weinberg, 2011). Most tumor types show phenotypic and genotypic heterogeneity attributable in large part to different mutational events accumulating throughout the existence of each tumor. The composite tumor mutation landscape is due to a combination of endogenous and exogenous sources with marked differences between cancer types. Knowledge of the intrinsic (bio)chemical preferences of mutagens together with more recent advances in DNA sequencing and bioinformatics have made it possible to break complex mutation landscapes into simpler individual components [reviewed by (Alexandrov and Stratton, 2014; Helleday et al., 2014; Roberts and Gordenin, 2014)]. In some cases, cause-and-effect relationships may be inferred because a given mutation spectrum or signature is only attributable to a single mutagen. The mutation landscape of melanoma, for instance, is often dominated by an ultra-violet (UV) light mutation signature, C-to-T mutations in dipyrimidine motifs due to UV-induced nucleobase crosslinking and adenine insertion opposite crosslinked lesions during DNA replication. In comparison, C-to-T mutations in CG dinucleotide motifs in many different cancer types are attributable to water-mediated deamination of methylated cytosines followed by DNA replication. The former process is avoidable and only accrues with UV light exposure, whereas the latter is unavoidable and accumulates stochastically over time [thus associated with biological age and often referred to as an aging mutation signature (Alexandrov et al., 2015)].

Another large source of C-to-T mutations in the genomes of many different tumor types is APOBEC3 (A3)-catalyzed DNA cytosine deamination (Alexandrov et al., 2013; Burns et al., 2013a; Burns et al., 2013b; Nik-Zainal et al., 2012; Roberts et al., 2013). Human cells have the potential to express up to seven different A3 family DNA cytosine deaminases (A3A-D, A3F-H). These enzymes function normally as an integral arm of the innate immune response to viral infections [reviewed by (Harris and Dudley, 2015; Malim and Bieniasz, 2012; Simon et al., 2015)]. However, A3-mediated deamination of genomic cytosines to uracils (C-to-U) can lead directly through DNA replication or indirectly through uracil excision repair and DNA replication to C-to-T mutations. In comparison to the aforementioned signatures, the primary APOBEC mutation signature is defined by C-to-T mutations in TC dinucleotides followed by adenine or thymine nucleobases to distinguish it from the aging signature (*i.e*., TCA and TCT motifs). A secondary APOBEC mutation signature is C-to-G transversion in the same trinucleotide motifs, which is likely due to REV1-catalyzed cytosine insertion opposite abasic lesions created by the combined action of cytosine deamination and uracil excision (Chan et al., 2013). The APOBEC mutation signature is evident in over half of all cancer types and, importantly, often dominates the mutation landscape of cervical, bladder, breast, head/neck, and lung tumors [reviewed by (Helleday et al., 2014; Roberts and Gordenin, 2014; Venkatesan et al., 2018){].

A major factor hindering broad appreciation of the APOBEC mutation process and efforts for clinical translation is a clear mechanistic determination of whether it is a causal driver or an inconsequential passenger in the overall tumor phenotype (drivers being potentially actionable and passengers being biomarkers at best). In support of a driver mechanism, first, both individual A3 enzymes as well as the overall APOBEC mutation signature have been associated with poor clinical outcomes including drug resistance and metastasis (Cescon et al., 2015; Chen et al., 2017; Glaser et al., 2018; Law et al., 2016; Sieuwerts et al., 2017; Sieuwerts et al., 2014; Walker et al., 2015; Xu et al., 2015; Yan et al., 2016). Second, a handful of *bona fide* driver mutations occur in APOBEC signature motifs, most prominently TCA to TTA mutations result in E542K and E545K changes in PIK3CA (Angus et al., 2019; Bertucci et al., 2019; Cannataro et al., 2019; Faden et al., 2017; Gillison et al., 2019; Henderson et al., 2014; Lefebvre et al., 2016). Third, high-risk human papillomavirus (HPV) infections cause upregulation of *A3A* and *A3B* gene expression, and HPV-positive tumors have stronger APOBEC mutation signatures than virus-negative tumors (Henderson et al., 2014; Mori et al., 2015; Mori et al., 2017; Vieira et al., 2014; Warren et al., 2015). In contrast and in support of a passenger model, first, A3A-mediated deamination of the single-stranded loop region of stem-loop structures (a mesoscale genomic feature) almost exclusively occurs in passenger genes (Buisson et al., 2019). Second, tumor genomes can contain hundreds to thousands of APOBEC signature mutations and almost all fail to recur between different patients implying little or no selective advantage (Angus et al., 2019; Bertucci et al., 2019; Nik-Zainal et al., 2016). Third, *A3A* and *A3B* expression is induced by interferon and inflammatory signals respectively, suggesting upregulation may be a consequence (not a cause) of other events in the overall tumor microenvironment (Koning et al., 2011; Leonard et al., 2015; Lucifora et al., 2014; Maruyama et al., 2016; Refsland et al., 2010; Siriwardena et al., 2019; Stenglein et al., 2010; Thielen et al., 2010). Additionally, the driver versus passenger question is complicated by a debate over which A3 enzyme or enzymes is the causal source of the APOBEC mutation signature in human cancer with A3A, A3B, and A3H as leading candidates [*e.g*., {Burns, 2013 #4434;Burns, 2013 #4657;Nik-Zainal, 2014 #5077;Caval, 2015 #5476;Caval, 2014 #5066;Middlebrooks, 2016 #6595;Chan, 2015 #5423;Cortez, 2019 #6646;de Bruin, 2014 #5124;Starrett, 2016 #5697;Yamazaki, 2019 #6656;Petljak, 2019 #6639;Serebrenik, 2019 #6641;Buisson, 2019 #6645}].

Here, two different models for tumorigenesis in mice are used to distinguish between driver and passenger roles for A3-catalyzed genomic DNA deamination in cancer. In the first murine tumor model, the Apc^Min^ model for colorectal carcinogenesis (Moser et al., 1990; Munteanu and Mastalier, 2014), transgenic expression of human A3A causes elevated frequencies of polyp formation and inflicts C-to-T mutations in APOBEC signature trinucleotide motifs. In the second model, the fumaryl-acetoacetate hydrolase (Fah) model for hepatocellular carcinoma (Keng et al., 2009; Wangensteen et al., 2008), all seven human A3 enzymes are tested individually including A3A, A3B, and A3H. However, only human A3A is able to significantly elevate hepatocellular tumor frequencies above baseline levels. In addition, A3A inflicts both primary (C-to-T) and secondary (C-to-G) mutations in APOBEC signature trinucleotide motifs indicating Rev1 functionality in hepatocytes during the transformation process. Additional bioinformatic analyses strongly indicate that the A3A mutation signatures induced in these murine cancer models closely resemble the overall APOBEC mutation signature in several different human tumors. These studies combine to demonstrate that A3A is capable of catalyzing mutation and actively driving carcinogenesis *in vivo* and that a proportion of the overall APOBEC mutation signature in human cancer is attributable to this enzyme.

## RESULTS

### Human A3 Expressing Mice

In contrast to humans with seven different A3 enzymes, mice encode only a single orthologous APOBEC3 protein (mA3) that manifests lower catalytic activity and localizes to the cytoplasmic compartment (MacMillan et al., 2013). Accordingly, APOBEC signature mutations have not been reported in any murine cancer model. Two strategies were used to create animals capable of expressing human APOBEC enzymes. The first was conditional expression of A3B from a multicopy transgene (Figure S1A-B). This approach appeared to work initially as evidenced by at least 2 generations of pups visibly expressing a linked DsRed marker (Figure S1C). However, both A3B functionality and DsRed expression were lost within 3 generations, potentially due to leaky deaminase expression from the transgene itself. Molecular analyses revealed that the *A3B* minigene was truncated and functionally inactivated and the linked *DsRed* cassette had accumulated 13 *de novo* base substitution mutations (Figure S1D-E).

The second approach was standard transgenesis, as reported initially in the context of antiviral studies (Stavrou et al., 2014). After 10 blastocyst injections and transgenesis attempts with a CAG promoter-driven *A3A* cDNA expression construct, two independent A3A lines were established: A3A^high^ with high levels of expression and activity and A3A^low^ with lower levels (Stavrou et al., 2014) (**Figure 1A**). However, similar to what occurred with the *A3B* transgenic mice, after multiple generations of breeding, we discovered through RNA sequencing that the A3A^high^ strain lost the C-terminal half of the *A3A* gene (Figure S2A). Sanger sequencing of cDNA generated from the A3A^high^ line revealed a deletion that fused *A3A* in-frame to the endogenous *Ngly1* gene (Figure S2B). The encoded chimeric protein is unlikely to be a functional DNA deaminase due to loss of essential A3A structural elements (Bohn et al., 2015; Kouno et al., 2017; Shi et al., 2017). A3A expression in the A3A^low^ animals was confirmed in multiple tissues, including the small intestine and colon by RT-PCR and RT-qPCR, as well as by a ssDNA deamination assay (**Figure 1B-D**). However, A3A^low^ expression was not high enough to detect with immunoblotting or immunohistochemistry using a custom mAb and validated protocols (Brown et al., 2019) (multiple attempts not shown). A3A^low^ expressing animals showed no overt phenotypes, normal life-spans, and normal Mendelian inheritance in 3 different animal facilities (Figure S3 and data not shown). These observations indicated that low levels of A3A are alone insufficient for tumorigenesis.

**Figure 1.**
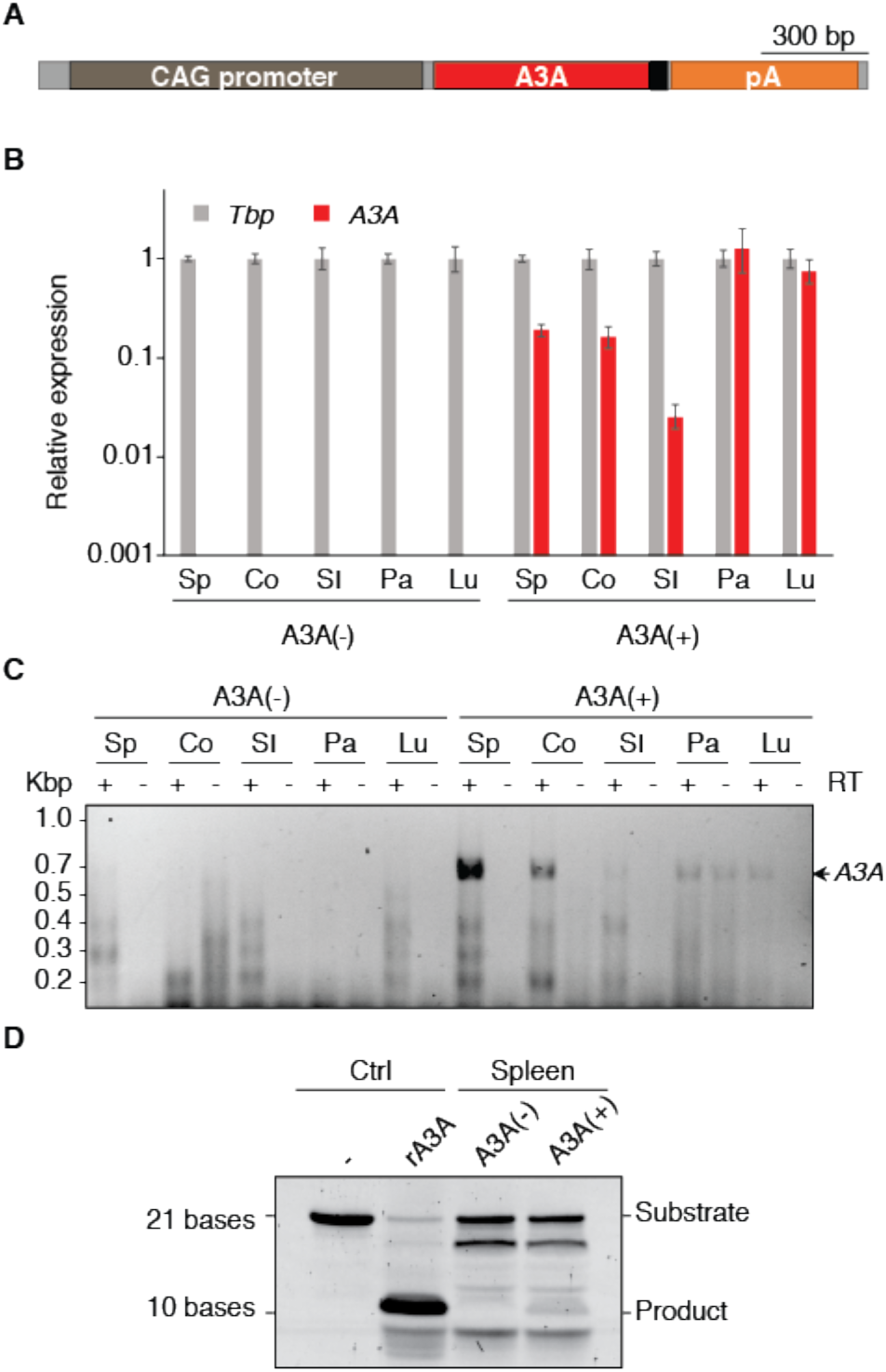
*A3A* expression and activity *in vivo*. **(A)** Transgene schematic with *A3A* in red and *myc* epitope in black. **(B)** RT-qPCR of *A3A* mRNA levels relative to those of the housekeeping gene *Tbp* (mean +/-SD for 3 technical replicates; Sp – spleen, Co – colon, SI – small intestine, Pa – pancreas, Lu – Lung). **(C)** Agarose gel image of full-length *A3A* cDNA amplified by RT-PCR from the indicated tissues. Parallel reactions excluding RT show minimal genomic DNA contamination (only pancreas). Arrow indicates size of *A3A* product. **(D)** Deamination activity in spleen extracts from A3A(-) and A3A(+) mice. Deamination substrate with no lysate or with 1 nM purified recombinant (r)A3A added to lysate are controls.

### Human A3A Promotes Intestinal Tumor Formation

At least one initial predisposing event is likely to be required for the APOBEC3 mutagenesis program to become active and influence tumor evolution (Olson et al., 2018; Siriwardena et al., 2016; Swanton et al., 2015; Venkatesan et al., 2018). Standard breeding was therefore done to combine A3A^low^ (hereafter A3A) with Apc^Min^, which is a well-established model for intestinal tumor development (Moser et al., 1990; Munteanu and Mastalier, 2014) (**Figure 2A**). This model was chosen due to high penetrance (all animals develop polyps), the quantifiable nature of polyp formation (dozens per animal), and a clearly defined analysis endpoint (sacrifice at 4 months of age prior to inevitable intestinal tract blockage and death). Polyp formation in A3A/Apc^Min^ and Apc^Min^ mice derived from the same crosses was analyzed at 4 months. In one series of experimental crosses, two of the A3A/Apc^Min^ animals died due to intestinal blockage prior to the 4-month endpoint. However, the remaining 12 animals made it to the endpoint for detailed histological and molecular analyses. The impact of A3A was immediately apparent with an approximate 1.5-fold increase in the median number of visible polyps in comparison to animals with Apc^Min^ alone (U=105.5, p=0.009 by two-sided Wilcoxon rank sum test; **Figure 2B-C**). Polyp formation was unevenly distributed through the intestinal tract with the greatest difference observed in the ileum (**Figure 2D**). Histopathologic examination of the polyps in the colon and small intestine revealed multiple, exophytic, pedunculated or sessile growths exhibiting adenomatous features, and neoplastic cells were characterized by columnar or cuboidal shape, ovoid, occasionally hyperchromatic nuclei, and formed tubular structures (**Figure 2E** and Figure S4). High-grade cytologic atypia with nuclear pleomorphism, increased mitotic figures (2-5 per high-power field) and tubular necrosis were observed in adenomas of both groups. The average size of the lesions did not appear to significantly differ between A3A/Apc^Min^ and Apc^Min^ mice. A second series of experimental crosses at a different animal facility yielded similar A3A-dependent increases in polyp formation and histological features (Figure S5).

**Figure 2.**
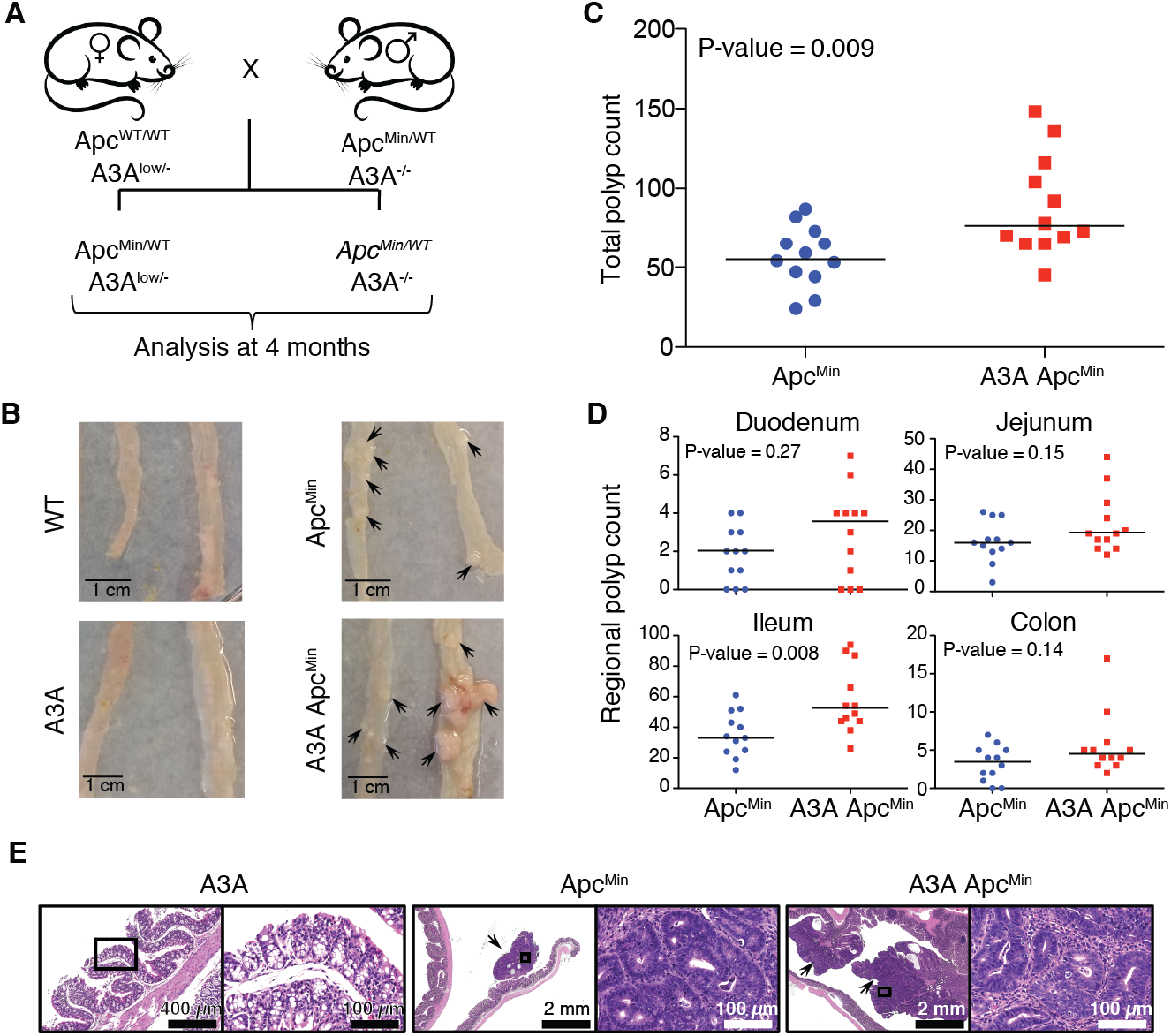
Human A3A promotes tumorigenesis in Apc^Min^ mice. **(A)** Breeding schematic for relevant groups. **(B)** Representative images of small intestine (left) and distal colon (right) from WT, A3A, Apc^Min^, and A3A/Apc^Min^ mice. Black arrowheads indicate polyp locations. **(C)** Total polyp counts in Apc^Min^ and A3A/Apc^Min^ mice (n=12; median shown by horizontal line). **(D)** Polyp counts for the indicated intestinal regions of Apc^Min^ and A3A/Apc^Min^ mice (subdivision of data in panel C). **(E)** Representative low- and high-power photomicrographs of indicated H&E stained colon sections. Image on the right side of each specimen is a high-power magnification of the region marked in a box in the left low-power image. Arrowheads point to polyps.

### Base Substitution Mutation Signatures in Human A3A Expressing Intestinal Tumors

DNA from colon polyps from A3A/Apc^Min^ and Apc^Min^ animals was subjected to whole exome sequencing, together with matched normal tail DNA. *A3A* RNA expression status in polyps was verified by RT-PCR (Figure S6). Single nucleotide variants (SNVs) present in the polyp exome profile, but absent in the matched normal DNA, were considered lesion-specific mutations. Interestingly, more mutations were detected on average in the A3A expressing tumors (t = −2.12, p = 0.037, one-sided two-sample t-test) (**Figure 3A**). Analysis of the different point mutation types showed that this increase in A3A expressing tumors is due largely to an elevated contribution of C-to-T mutations in non-CG motifs (**Figure 3B**). Evaluation of the trinucleotide contexts revealed that A3A expressing polyps have increased C-to-T mutations within TC motifs (**Figure 3C-D**).

**Figure 3.**
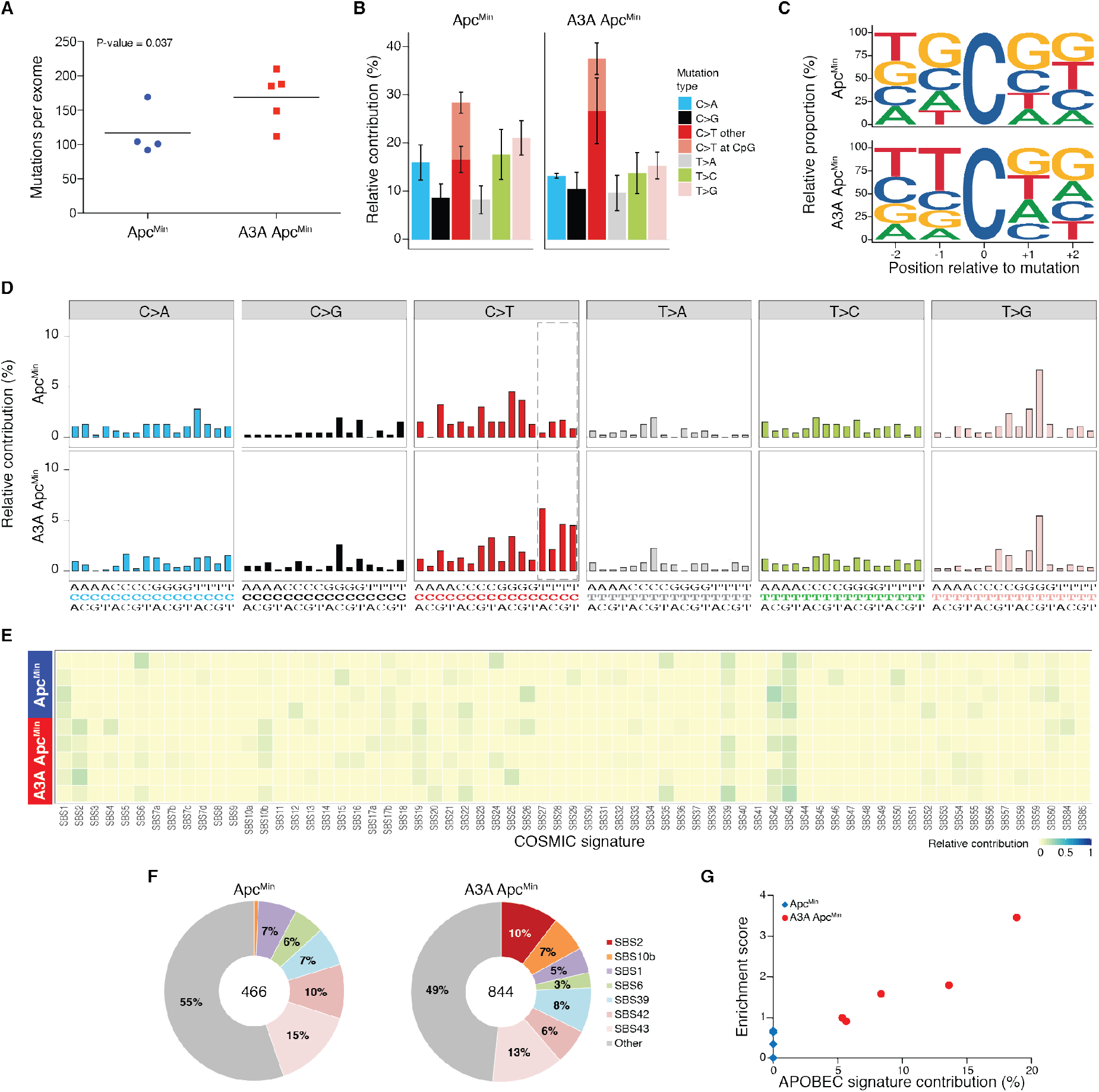
APOBEC signature mutations in polyps from human A3A expressing animals. **(A)** Dot plot of total base substitution mutations per tumor exome in Apc^Min^ versus A3A/Apc^Min^ animals (n=4 and 5, respectively; mean shown by horizontal line). **(B)** Barplot representing relative contribution of the 6 different types of base substitutions to the total mutation burden (mean +/-SD). C-to-T transitions further categorized as occurring within CG dinucleotides or not. **(C)** Sequence logos for all C-to-T mutations in Apc^Min^ versus A3A/Apc^Min^ tumors. **(D)** Barplots representing composite trinucleotide mutation profiles for all base substitutions in Apc^Min^ versus A3A/Apc^Min^ tumors. Dotted box indicates location of canonical A3-preferred trinucleotides. **(E-F)** Heatmap and pie charts summarizing the relative contribution of each mutation signature to the observed constellation of mutations in Apc^Min^ versus A3A/Apc^Min^ tumors. SBS2 (APOBEC) is only evident in A3A expressing tumors and SBS10b is enriched in A3A-expressing tumors. **(G)** Scatterplot of APOBEC signature contribution to TCW motif enrichment score.

The trinucleotide mutational spectra of each tumor were deconstructed in relation to the current list of COSMIC mutational signatures (v3 - bioRxiv preprint doi: http://dx.doi.org/10.1101/322859). Strikingly, there was no contribution whatsoever of APOBEC-related single base substitution signatures (SBS2 and SBS13) in any of the control tumors, whereas each of the A3A-expressing tumors showed clear evidence for SBS2 (APOBEC), ranging from 5%-19% of total mutations, though no SBS13 was observed (**Figure 3E-G** and Figure S7). Intriguingly, one additional signature (SBS10b) was also only evident in A3A-expressing tumors, suggesting that this may be a linked downstream mutational outcome (**Figure 3E-F**). Enrichment scores of mutations within TCW motifs were significantly elevated in A3A-expressing tumors in relation to control tumors (t=-2.73, p =0.043, two-sided two-sample t-test) and also strongly associated with APOBEC signature contribution (**Figure 3G**). The aging signature (SBS1) was not strong in any of the polyps, likely due to the relatively young age of the animals. In support of APOBEC signature being dominant in the A3A/Apc^Min^ tumors, a *de novo* deconstruction of the trinucleotide mutation data using non-negative matrix factorization (Alexandrov et al., 2013) also led to the generation of a pattern strongly resembling SBS2 (APOBEC) in the A3A/Apc^Min^ tumors but not the APC^Min^ tumors (Figure S8).

### Human A3A May Also Trigger Chromosomal Aberrations Including *Apc* Loss-of-Heterozygosity (LOH)

Mutations predicted to cause changes in coding sequences were further analyzed, revealing that 21-30% of point mutations detected in each tumor were non-synonymous. Whereas a range of different genes were mutated in these tumors, known adenocarcinoma drivers (Munteanu and Mastalier, 2014) such as *Tp53*, *Pten*, *K-Ras*, *Tgfbr2*, and *Smad4* were unchanged. This is consistent with the pathological classification of the polyps as adenomas that have yet to undergo invasive transformation. However, for both Apc^Min^ and A3A/Apc^Min^, all of the sequenced tumors showed LOH for the chromosome 18 region spanning the *Apc*^*WT*^ gene leading to Apc loss of function (Figure S9). Because no other recurrent alterations distinguished the A3A/Apc^Min^ and Apc^Min^ groups and no LOH events were evident in the tail DNA from either group of animals, the increased polyp number may be due to a subset of A3A-catalyzed uracil lesions being processed into DNA breaks and contributing to higher rates of chromosomal instability including loss-of-heterozygosity.

### Human A3A Promotes Liver Carcinogenesis

We next turned to the Fah model for liver regeneration and hepatocellular carcinoma (Grompe et al., 1998; Keng et al., 2009; Wangensteen et al., 2008). *Fah*-null animals are defective in tyrosine metabolism and liver failure results from the toxic build-up of the primary Fah substrate (fumaryl-acetoacetate) and its precursor (maleyl-acetoacetate). However, if supplemented with the drug 2-(2-nitro-4-trifluoromethylbenzoyl)-1,3-cyclohexanedione (NTBC), which inhibits an enzyme upstream in the same anabolic pathway, these toxic metabolites fail to accumulate and the liver functions normally. The essential nature of the Fah enzyme and the regenerative capacity of the liver enables genetic complementation of the deficiency upon delivery of a functional *Fah* transgene and withdrawal of NTBC to select for liver repopulation by complemented cells (Grompe et al., 1998). We therefore leveraged these properties and constructed a set of *Fah* complementation vectors for delivery by hydrodynamic transfer and heritable transmission by *Sleeping Beauty* (*SB*)-catalyzed integration into genomic DNA (schematic in **Figure 4A**). The primary *SB* payload has a PGK promoter driving *Fah* expression and a CAG promoter for direct expression of individual human A3 family members and IRES-mediated expression of luciferase. A secondary *SB* payload expresses a short hairpin RNA against *Tp53*, which promotes tumor development (Keng et al., 2009; Wangensteen et al., 2008). These genetic payloads are delivered simultaneously to the genomes of hepatocytes through SB11-mediated transposition (Keng et al., 2009; Wangensteen et al., 2008).

**Figure 4.**
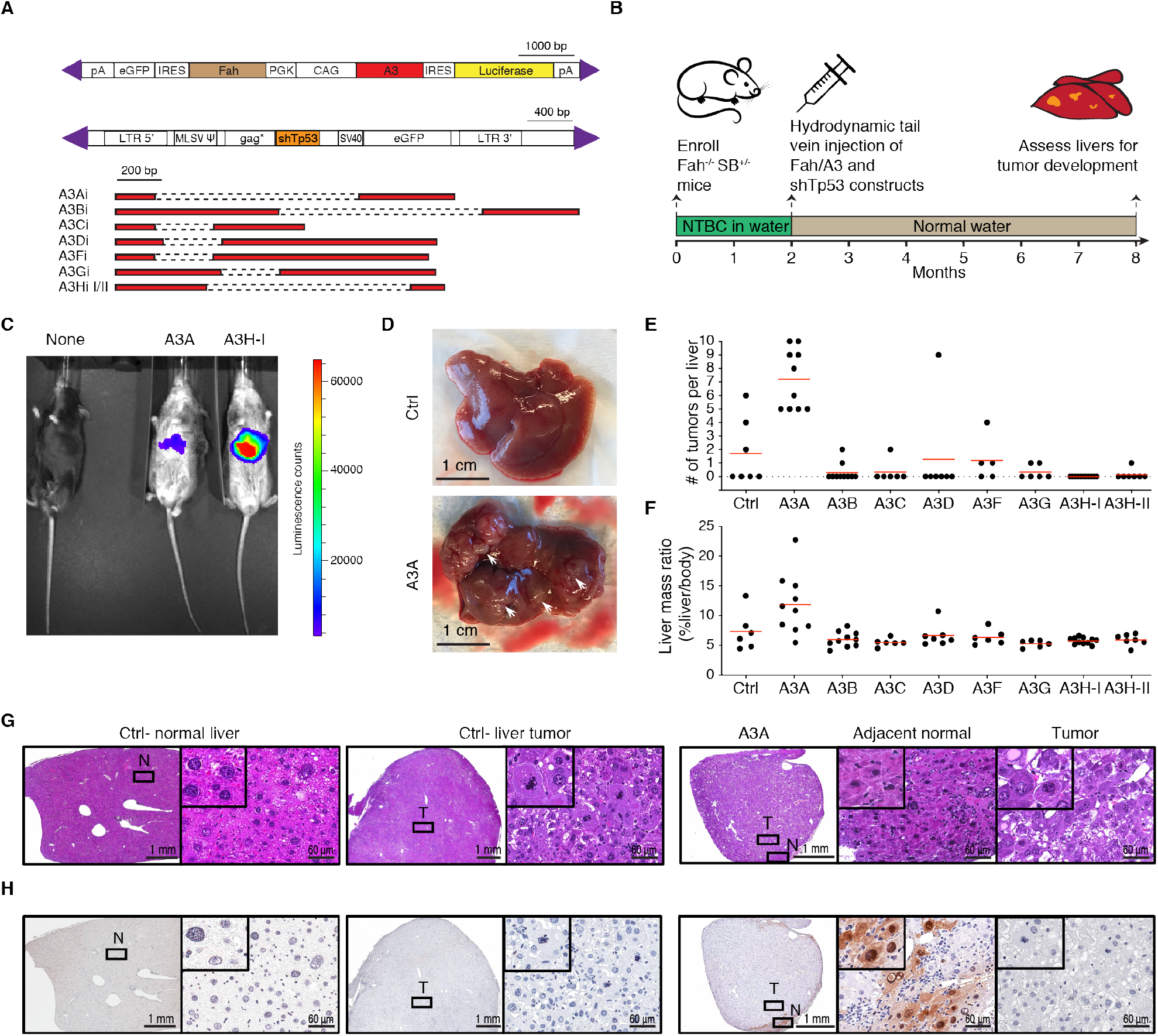
A3A causes hepatocellular carcinogenesis in Fah liver regeneration model. **(A)** Schematics of *Fah*-*A3(A-D, F-H)* and sh*Tp53* expressing constructs injected into *Fah*-null SB11 animals. Purple triangles indicate IR/DR signal sequences required for transposition by SB11. Human *A3* minigene schematics are shown below with dashed lines representing introns. **(B)** Overview of experimental workflow. **(C)** Luminescence of representative 7.5 month-old liver regenerated mice. **(D)** Representative images of control and A3A Fah repopulated livers. White arrows point to tumors. **(E-F)** Tumor burdens and liver/body mass ratios for the indicated groups. **(G)** Representative photomicrographs of H&E-stained sections of non-neoplastic (normal - N) and tumorous (T) livers originating from control and A3A Fah animals. Images on the right represent high-power magnification(s) of the regions marked in a box in the left low-power images. Inset images are magnified an additional 4-fold. **(H)** IHC staining of the same tissues as panel (G) with a rabbit α-human A3A/B/G mAb (5210-87-13). Layout descriptions are the same as panel (G).

The major strength of the Fah system is that it enables simultaneous testing of the 7 different human A3 enzymes in parallel in an otherwise isogenic system *in vivo* (workflow in **Figure 4B**). *Fah*-null SB11 animals were enrolled at birth and supplemented with NTBC drinking water and then, at 2 months age, provided with normal drinking water and randomized for hydrodynamic injections with the sh*Tp53* vector and a control Fah vector (no A3) or an *A3*-expressing Fah vector. Representative animals expressed luciferase in liver tissues in comparison to non-injected *Fah*-null SB11 animals (**Figure 4C**). At 8 months of age (6 months after hydrodynamic delivery), the entire cohort was sacrificed for comprehensive pathological and molecular analyses. The most visually striking result was a uniformly high tumor number in the livers of A3A-expressing animals (range = 5-10; mean = 7.2 and p<0.0001 by one-way ANOVA; **Figure 4D-E**). Both control (non-A3) and all other A3-expressing animals had none or 1 visible tumor per liver with occasional outliers with more events (mean tumor numbers not statistically different from control group; p-value range = 0.18 – 0.99 by one-way ANOVA). Accordingly, livers in A3A-expressing group were an average of nearly 100% larger than those from the control or other A3-expressing groups (average liver mass ratio of 0.12 versus an average range of 0.06-0.07; p = 0.013 by one-way ANOVA; **Figure 4F**). The heritable integration and integrity of all *A3* and sh*Tp53* expression constructs was confirmed by diagnostic PCR assays of liver tissue genomic DNA (Figure S10A). Moreover, RTqPCR assays confirmed expression of the various *A3* constructs at the RNA level (Figure S10B), and immunoblots were used to show expression of A3A, A3B, A3C, A3G, and A3H haplotypes I and II at the protein level (Figure S10C). Furthermore, single-stranded DNA deaminase activity assays were used to demonstrate activity of A3A and A3B in liver whole cell extracts (Figure S10D). These results combined to indicate that only human A3A has the capacity to drive tumor formation in the murine Fah system.

Fah tumor pathology and immunohistology were also informative. First, hematoxylin and eosin (H&E) stained liver sections from A3A-expressing animals revealed tumorous growths with lobular architecture and marked histologic heterogeneity in the neoplastic cell population. The cancer cells primarily featured a solid and/or trabecular growth pattern and showed size and shape variation, pronounced cytologic atypia, nuclear pleomorphism, hyperchromatism, and increased numbers of mitotic figures. In addition, focal areas of necrosis, intracellular edema, apoptotic phenomena, and mild-to-moderate lymphocytic inflammation were seen (**Figure 4G**). Second, attempts to detect A3A at the protein level using a custom mAb and a validated IHC assay (Brown et al., 2019) revealed an additional layer of heterogeneity. In most instances, the bulk of the tumor itself showed limited/no evidence for A3A protein expression, whereas residual, adjacent to the tumor, non-neoplastic liver parenchyma often displayed aggregates of hepatocytes with moderate to strong nuclear and cytoplasmic staining (**Figure 4H**). These results are consistent with prior reports for cell-wide human A3A localization (Caval et al., 2015; Chen et al., 2006; Hultquist et al., 2011; Land et al., 2013)} and its capacity to exert cytotoxicity (Buisson et al., 2017; Caval et al., 2015; Landry et al., 2011; Stenglein et al., 2010)}. One possible interpretation of this remarkable heterogeneity is that, despite a strong positive selective pressure for Fah function, an even stronger selection against A3A may be occurring during tumorigenesis that results in A3A inactivation (**Discussion**).

### Base Substitution Mutation Signatures in Liver Tumors Attributable to A3A

Whole exome sequencing was done for 8 liver tumors from 4 separate animals, as well as normal tissue (tail) from each animal to enable unambiguous mutation calls. Mutation numbers ranged widely, even between tumors from the same animal, consistent with independent transformation events (n = 127 to 4082) (**Figure 5A**). The majority of these base substitutions are C-to-T transitions in TC motifs preceded by a C or T at the −2 position and followed by an A or T at the +1 position (**Figure 5B-D**). As expected, these tumors have a strong SBS2 (range = 0 - 46%); however, unlike the A3A Apc^Min^ tumors above, several also show a clear SBS13 (range = 0 - 13%; **Figure 5E-G** and Figure S11-12). SBS7a and, as above, SBS10b are only evident in tumors with strong A3A mutation signatures suggesting possible mechanistic linkage (*e.g*., SBS7a may be due to promiscuous A3A-catalyzed deamination of less preferred TCC contexts; **Figure 5F**). Finally, APOBEC mutation signatures SBS2 and SBS13 correlate positively with enrichment scores for cytosine mutation at TCW relative to other cytosine contexts (Spearman’s rank correlation = 0.96, p = 0.0002 and Spearman’s rank correlation = 0.91, p = 0.0016, respectively; **Figure 5G-H**). These results demonstrate that A3A is capable of causing both APOBEC signatures SBS2 and SBS13 *in vivo*.

**Figure 5.**
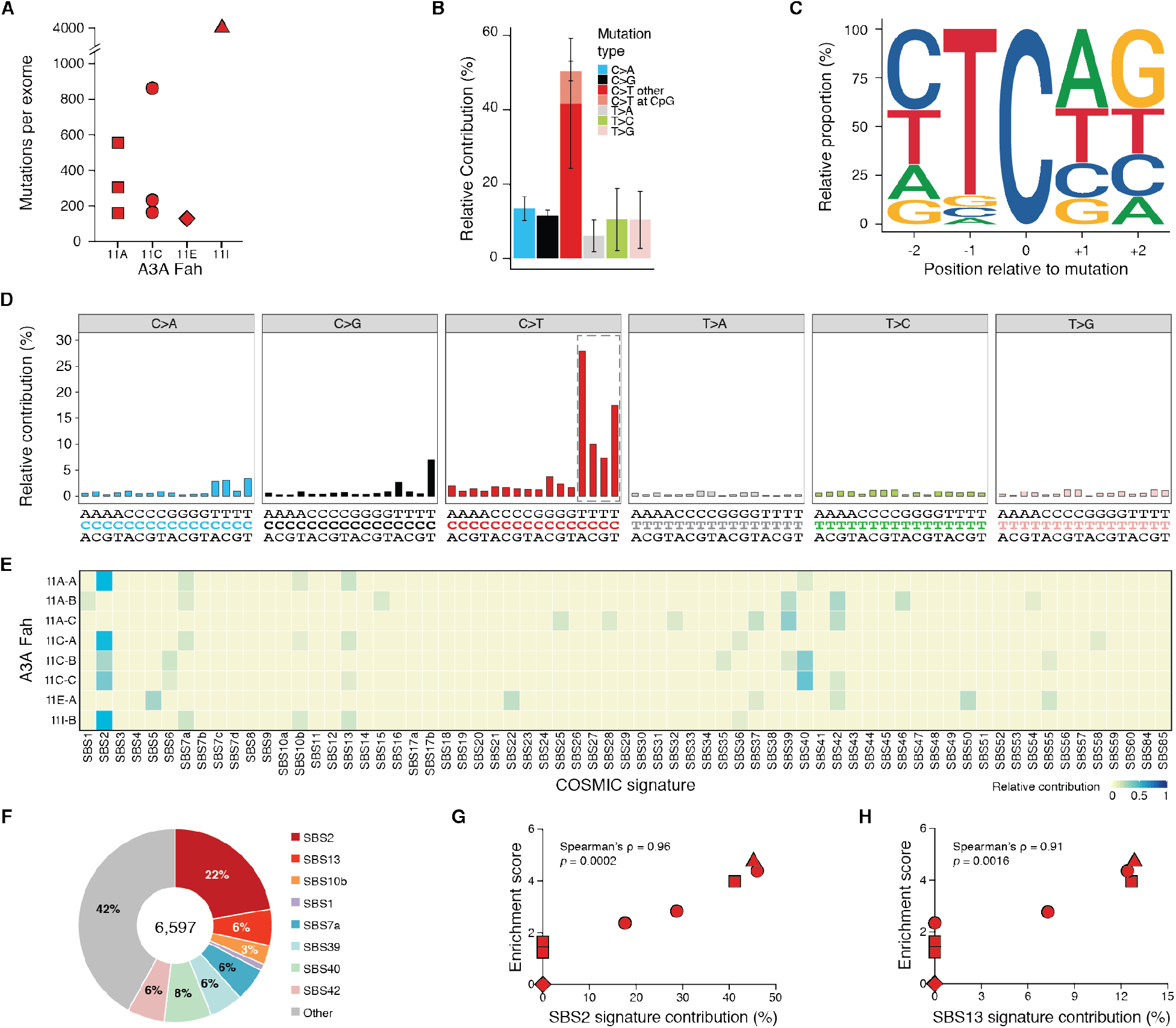
APOBEC signature mutations in liver tumors from A3A Fah mice. **(A)** Dot plot of total base substitution mutations per tumor exome in A3A Fah animals. Different symbol shapes report data from distinct animals (n=8 tumors from 4 independent animals). **(B)** Barplot showing the percentage of the 6 different types of base substitutions in A3A Fah tumors (mean +/-SD). C-to-T transitions further categorized as occurring within CG dinucleotides or not. **(C)** Composite sequence logo for all C-to-T mutations occurring in A3A Fah tumor exomes. **(D)** Barplots representing composite trinucleotide mutation profiles for all base substitutions in A3A Fah tumors. Dotted box indicates location of canonical A3-preferred trinucleotides. **(E-F)** Heatmap and pie charts summarizing the relative contribution of each mutation signature to the observed constellation of mutations in A3A Fah tumors. Signatures SBS2 and SBS13 (APOBEC) predominate over other signatures in A3A Fah tumor exome data. **(G-H)** Scatterplots of TCW enrichment scores versus SBS2 and SBS13 contributions, respectively. Different symbol shapes denote distinct animals of origin (matching those in A).

### Murine and Human Tumor APOBEC Mutation Signature Comparisons

To compare the APOBEC mutation signatures observed in murine A3A tumor models and those occurring in human cancers, an unbiased Euclidean distance dendrogram was generated using the nucleobases surrounding the mutated cytosines in our murine colon and liver tumor data sets (above), TCGA human cancer data sets (Cancer Genome Atlas Research Consortium, 2013), and previously reported data for A3A and A3B mutagenesis in yeast (Chan et al., 2015) (**Figure 6**). To minimize potential artifacts as well as overlapping signals from other mutational processes including aging, these analyses utilized all C-to-T mutations in NTC(A/C/T) tetranucleotide motifs and excluded those in CG motifs. The first notable point is that the A3A-induced signature in both the Apc^Min^ and Fah tumors is YTCW and clustered very tightly at the top of the tree. Second, the A3A-induced signature in Apc^Min^ and Fah tumors clusters adjacent to the tetranucleotide C-to-T mutation signatures of the largest APOBEC signature cancer types in humans – cervical, bladder, HPV-positive head/neck, and lung adenocarcinoma. Third, the tetranucleotide C-to-T mutation signatures from *A3B*-null breast cancers and A3A-expressing yeast cluster closely and as an outgroup of the aforementioned signatures. Fourth, the tetranucleotide C-to-T mutation signatures from A3B-expressing yeast cluster closely to those of human kidney and adrenocortical tumors but still within the APOBEC dominated half of the dendrogram. Last, the tetranucleotide C-to-T mutation signatures from all other human tumor types cluster in various ways in the other half, largely non-APOBEC, of the dendrogram. These analyses are consistent with the idea that A3A is a driver of mutation in multiple human tumor types including those of the cervix, bladder, HPV-positive head/neck, and lung.

**Figure 6.**
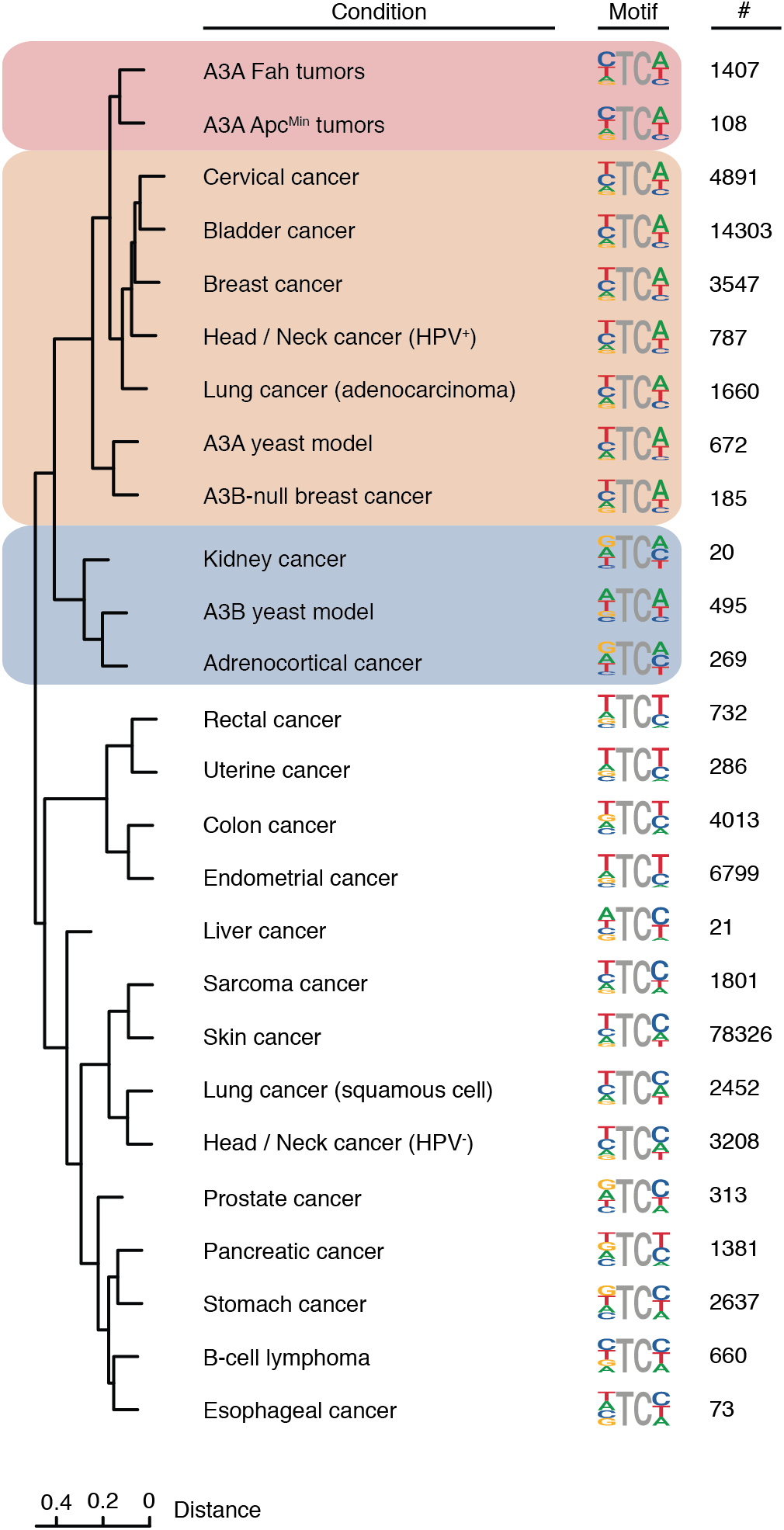
APOBEC mutation signature comparisons for murine and human tumors. Euclidian distance dendrogram depicting hierarchical relationships between C-to-T mutation signatures of human A3A in mouse tumors, human A3A and A3B in yeast, and the indicated human cancers. Each tetranucleotide signature is shown next to the relevant branchtip along with the number of mutations contributing to each analysis. See text for additional details.

## DISCUSSION

The studies described here were initiated to determine whether the APOBEC mutation process is a driver or a passenger in the overall tumorigenesis process. Several lines of prior evidence support both models and, importantly, none has been definitive (see **Introduction**). Here we show that human A3A is able to drive tumorigenesis in two different murine systems. Low, constitutive levels of A3A from a stable transgene promote polyp formation in the Apc^Min^ model for colorectal cancer. Higher, likely transient levels of A3A from hydrodynamic delivery of a SB-transposable element cause hepatocellular carcinogenesis in a *Tp53*-depleted Fah liver regeneration system. In both systems, A3A inflicts a hallmark APOBEC mutation signature in the genomes of tumors. These are the first demonstrations *in vivo* in a complex mammalian system that a human A3 enzyme catalyzes mutagenesis and promotes tumorigenesis. Moreover, the strong similarity of the A3A mutation signature in these murine tumor models and the overall APOBEC mutation signature in cervical, bladder, HPV-positive head/neck, and lung cancers provides additional evidence for an active role of this DNA deaminase in human tumor development and evolution.

None of the other human A3 enzymes, including A3B, showed an ability to accelerate tumor development in the *Tp53*-depleted Fah liver regeneration system (and testing the entire family in the Apc^Min^ system is not possible due to lack of transgenics for each human enzyme). There are several possible explanations for this negative result. One factor may be the fixed 6-month timeframe from hydrodynamic delivery to analysis. A second consideration is the fact that A3A is more active biochemically than the next most potent human A3 enzymes A3B and A3H (Adolph et al., 2017; Buisson et al., 2019; Carpenter et al., 2012; Cortez et al., 2019; Ito et al., 2017). A third consideration could be negative interference by an as yet unidentified factor. For instance, both intra/inter-A3 oligomerization and RNA binding are known to inhibit the activity of multiple A3 family DNA deaminases including A3B and A3H (Cortez et al., 2019; Shaban et al., 2018). However, these considerations are tempered by the fact that our original human A3B transgenic line was lost early in these studies likely due to its exerting negative selective pressure and catalyzing self-inactivation. A fourth consideration is the possibility of anti-human A3 immune responses, which may be lower for A3A in comparison to other family members.

Indeed, it will be very important to address the potential role of immune responses in future studies. The majority of liver tumors in the Fah regeneration model showed A3A mutation signatures but little to no detectable intra-tumor A3A staining by IHC. For instance, the tumor with the highest mutation load in **Figure 5A** (4082 base substitutions) stained IHC-negative for A3A in **Figure 4G-H**, despite adjacent hepatocellular parenchyma showing A3A expression cell-wide. This apparent incongruity strongly suggests a dynamic interplay between positive selection for restoration of Fah function, negative selection against A3A as a genotoxic and potentially cytotoxic stress (further complicated by tight linkage of *A3A* and *Fah* on the same transposable element), and additional negative selective pressure by immune responses against the foreign proteins encoded by the transposon constructs. Another related possibility is immune responses against foreign antigens (neoantigens) created by A3A-catalyzed mutation. Additional studies will be needed to deconvolute these competing pressures and expand the utility of the systems described here for studying the APOBEC mutation program in cancer.

It is curious that the A3A Apc^Min^ colon tumors only manifest C-to-T transitions in TCA/T motifs (SBS2), whereas the A3A Fah liver tumors additionally show C-to-G transversions in the same preferred motifs (SBS13). This may be due simply to the lower mutation loads observed in the Apc^Min^ system, and an analysis of larger tumor numbers or whole genome sequences may reveal evidence for SBS13. Alternatively, this difference may be due to lower rates of uracil excision in the intestinal crypt cells that ultimately give rise to polyps in the Apc^Min^ model. Additionally, these differences in mutation signature may be due to low (or no) Rev1 translesion DNA synthesis in colon tissue, which is the only DNA polymerase known to insert cytosine nucleobases opposite abasic sites (Lawrence, 2002). In support, Rev1 overexpression was necessary to stimulate polyp formation and enhance mutagenesis in an MNU-dependent murine intestinal adenoma model (Sasatani et al., 2017).

Classic studies overexpressed distantly related deaminases *in vivo*, rat Apobec1 and murine Aicda (activation-induced deaminase), and reported hepatocellular carcinomas and T cell lymphomas and lung adenomas, respectively (Okazaki et al., 2003; Yamanaka et al., 1995). The liver tumors induced by rat APOBEC1 were attributed originally to dysregulated RNA editing and, as far as we are aware, chromosomal DNA mutations were not considered a possibility. Segments of select genes, *TCRβ* and *c-myc*, were sequenced in a subset of Aicda lymphomas and a bias toward C/G-to-T/A mutations was reported. Aicda transgenic animals also displayed shorter lifespans with increasing generation times, perhaps by mechanisms similar to the loss of the A3A^high^ and A3B transgenics described here. Similarly, AICDA (more commonly called AID) is implicated in carcinogenesis in specific human tissues, notably B cell and myeloid cell lymphomas and gastric cancers [reviewed by (Marusawa and Chiba, 2010; Robbiani and Nussenzweig, 2013)]. However, unlike AID, the human A3 enzymes are broadly implicated in carcinogenesis and arguably the second largest source of mutation across cancer. Thus, the A3A tumor models described here provide long-awaited *in vivo* demonstrations that this human DNA deaminase can indeed drive tumor development and is not simply a passenger mutation process.

It should be noted that mutational processes such as A3A/AID-catalyzed DNA cytosine deamination do not precisely fit the classical cancer definitions of “oncoprotein” and “driver”. These definitions may need to be expanded to include subclasses of tumor-driving processes, exemplified by “enablers” such as A3A/AID, that provide major sources of mutational fuel for tumor development and evolution. Although many exogenous and endogenous mutational processes contribute to the overall mutation landscape in human cancer, only a small number of processes such as A3A/AID are likely to be large and dominant enough to enable measurable changes in rates of tumorigenesis and trajectories of tumor development. Such enablers have the potential to provide large proportions of the mutational fuel for many different tumor types, which coupled with selective pressures for/against tumor development, result in the massive heterogeneity reported in the genomic DNA sequences of different human tumor types (as well as between individual tumors within distinct tumor types). It is therefore important to determine *in vivo*, as here for A3A, whether a particular source of mutation is strong enough to be defined as both a source of mutation and a driver of tumorigenesis. Such information will be critical to be able to fully investigate future diagnostic, prognostic, and therapeutic options with the ultimate goal of preventing or treating mutational drive.

## ACKNOWLEDGMENTS

We thank Hilde Nilsen and Douglas Yee for thoughtful feedback, Bobbi Tschida for training on hydrodynamic injections, Brian Dunnette for sharing expertise regarding the use of the Aperio ScanScope XT, and Kristin Blouch and David Ryan for assistance with mouse breeding. These studies were supported in part by grants from the National Cancer Institute (P01CA234228 to R.S.H.), the National Institute for Allergy and Infectious Disease (R01AI085015 to S.R.R.), National Institute for Aging (R21AG047114 to S.R.R.), the University of Minnesota College of Biological Sciences and Academic Health Center (to R.S.H.), the Randy Shaver Cancer Research and Community Fund (to R.S.H.), the Masonic Cancer Center Translational Working Group (to T.K.S.), and the Mezin-Koats Colorectal Cancer Research Fund grants (to T.K.S.). NIH training grants and career development awards provided salary support for R.L.-K. (T32HL007062 and F32CA232458-01) and M.C.J (T32CA009138 and F31CA243306). R.L.-K. is an Awardee of the Weizmann Institute of Science - National Postdoctoral Award Program for Advancing Women in Science. M.B.B. was supported in part by a Department of Defense Breast Cancer Research Program Predoctoral Fellowship (BC101124). R.S.H is the Margaret Harvey Schering Land Grant Chair for Cancer Research, a Distinguished University McKnight Professor, and an Investigator of the Howard Hughes Medical Institute.

## AUTHOR CONTRIBUTIONS

R.S.H. and S.R.R. conceptualized and supervised the overall project. E.K.L, R.L.-K., M.C.J., H.K., P.P.A., M.A.C., L.K.L., M.B.B., S.S., A.N.A., and T.K.S. performed experiments. R.L.-K. and H.K. performed validation studies. E.K.L., R.L.-K., M.C.J., G.J.S., and R.I.-V. did formal data analysis. E.K.L, R.L.-K., M.C.J., H.K., M.A.C., S.W., D.A.L., and T.K.S. contributed methodology. R.L.-K., M.C.J., M.B.B., D.A.L., T.K.S., S.R.R., and R.S.H. contributed to funding acquisition. R.L.-K., M.C.J., E.K.L., P.P.A., S.R.R., and R.S.H. drafted the manuscript and all authors contributed to revisions.

## REFERENCES

Adolph, M.B., Love, R.P., Feng, Y., and Chelico, L. (2017). Enzyme cycling contributes to efficient induction of genome mutagenesis by the cytidine deaminase APOBEC3B. Nucleic Acids Res 45, 11925–11940.

Alexandrov, L.B., Jones, P.H., Wedge, D.C., Sale, J.E., Campbell, P.J., Nik-Zainal, S., and Stratton, M.R. (2015). Clock-like mutational processes in human somatic cells. Nat Genet 47, 1402–1407.

Alexandrov, L.B., Nik-Zainal, S., Wedge, D.C., Aparicio, S.A., Behjati, S., Biankin, A.V., Bignell, G.R., Bolli, N., Borg, A., Borresen-Dale, A.L., et al. (2013). Signatures of mutational processes in human cancer. Nature 500, 415–421.

Alexandrov, L.B., and Stratton, M.R. (2014). Mutational signatures: the patterns of somatic mutations hidden in cancer genomes. Curr Opin Genet Dev 24, 52–60.

Angus, L., Smid, M., Wilting, S.M., van Riet, J., Van Hoeck, A., Nguyen, L., Nik-Zainal, S., Steenbruggen, T.G., Tjan-Heijnen, V.C.G., Labots, M., et al. (2019). The genomic landscape of metastatic breast cancer highlights changes in mutation and signature frequencies. Nat Genet.

Bertucci, F., Ng, C.K.Y., Patsouris, A., Droin, N., Piscuoglio, S., Carbuccia, N., Soria, J.C., Dien, A.T., Adnani, Y., Kamal, M., et al. (2019). Genomic characterization of metastatic breast cancers. Nature 569, 560–564.

Blokzijl, F., Janssen, R., van Boxtel, R., and Cuppen, E. (2018). MutationalPatterns: comprehensive genome-wide analysis of mutational processes. Genome medicine 10, 33.

Bohn, M.F., Shandilya, S.M., Silvas, T.V., Nalivaika, E.A., Kouno, T., Kelch, B.A., Ryder, S.P., Kurt-Yilmaz, N., Somasundaran, M., and Schiffer, C.A. (2015). The ssDNA mutator APOBEC3A is regulated by cooperative dimerization. Structure 23, 903–911.

Brown, W.L., Law, E.K., Argyris, P.P., Carpenter, M.A., Levin-Klein, R., Ranum, A.N., Molan, A.M., Forster, C.L., Anderson, B.D., Lackey, L., et al. (2019). A rabbit monoclonal antibody against the antiviral and cancer genomic DNA mutating enzyme APOBEC3B. Antibodies (Basel) 8.

Buisson, R., Langenbucher, A., Bowen, D., Kwan, E.E., Benes, C.H., Zou, L., and Lawrence, M.S. (2019). Passenger hotspot mutations in cancer driven by APOBEC3A and mesoscale genomic features. Science 364.

Buisson, R., Lawrence, M.S., Benes, C.H., and Zou, L. (2017). APOBEC3A and APOBEC3B activities render cancer cells susceptible to ATR inhibition. Cancer Res 77, 4567–4578.

Burns, M.B., Lackey, L., Carpenter, M.A., Rathore, A., Land, A.M., Leonard, B., Refsland, E.W., Kotandeniya, D., Tretyakova, N., Nikas, J.B., et al. (2013a). APOBEC3B is an enzymatic source of mutation in breast cancer. Nature 494, 366–370.

Burns, M.B., Temiz, N.A., and Harris, R.S. (2013b). Evidence for APOBEC3B mutagenesis in multiple human cancers. Nat Genet 45, 977–983.

Cancer Genome Atlas Research Network (2013). The Cancer Genome Atlas Pan-Cancer analysis project. Nat Genet 45, 1113–1120.

Cannataro, V.L., Gaffney, S.G., Sasaki, T., Issaeva, N., Grewal, N.K.S., Grandis, J.R., Yarbrough, W.G., Burtness, B., Anderson, K.S., and Townsend, J.P. (2019). APOBEC-induced mutations and their cancer effect size in head and neck squamous cell carcinoma. Oncogene.

Carpenter, M.A., Li, M., Rathore, A., Lackey, L., Law, E.K., Land, A.M., Leonard, B., Shandilya, S.M., Bohn, M.F., Schiffer, C.A., et al. (2012). Methylcytosine and normal cytosine deamination by the foreign DNA restriction enzyme APOBEC3A. J Biol Chem 287, 34801–34808.

Caval, V., Bouzidi, M.S., Suspene, R., Laude, H., Dumargne, M.C., Bashamboo, A., Krey, T., Vartanian, J.P., and Wain-Hobson, S. (2015). Molecular basis of the attenuated phenotype of human APOBEC3B DNA mutator enzyme. Nucleic Acids Res 43, 9340–9349.

Cescon, D.W., Haibe-Kains, B., and Mak, T.W. (2015). APOBEC3B expression in breast cancer reflects cellular proliferation, while a deletion polymorphism is associated with immune activation. Proc Natl Acad Sci U S A 112, 2841–2846.

Chan, K., Resnick, M.A., and Gordenin, D.A. (2013). The choice of nucleotide inserted opposite abasic sites formed within chromosomal DNA reveals the polymerase activities participating in translesion DNA synthesis. DNA Repair (Amst) 12, 878–889.

Chan, K., Roberts, S.A., Klimczak, L.J., Sterling, J.F., Saini, N., Malc, E.P., Kim, J., Kwiatkowski, D.J., Fargo, D.C., Mieczkowski, P.A., et al. (2015). An APOBEC3A hypermutation signature is distinguishable from the signature of background mutagenesis by APOBEC3B in human cancers. Nat Genet 47, 1067–1072.

Chen, H., Lilley, C.E., Yu, Q., Lee, D.V., Chou, J., Narvaiza, I., Landau, N.R., and Weitzman, M.D. (2006). APOBEC3A is a potent inhibitor of adeno-associated virus and retrotransposons. Curr Biol 16, 480–485.

Chen, T.W., Lee, C.C., Liu, H., Wu, C.S., Pickering, C.R., Huang, P.J., Wang, J., Chang, I.Y., Yeh, Y.M., Chen, C.D., et al. (2017). APOBEC3A is an oral cancer prognostic biomarker in Taiwanese carriers of an APOBEC deletion polymorphism. Nat Commun 8, 465.

Cortez, L.M., Brown, A.L., Dennis, M.A., Collins, C.D., Brown, A.J., Mitchell, D., Mertz, T.M., and Roberts, S.A. (2019). APOBEC3A is a prominent cytidine deaminase in breast cancer. PLoS Genet 15, e1008545.

Faden, D.L., Thomas, S., Cantalupo, P.G., Agrawal, N., Myers, J., and DeRisi, J. (2017). Multi-modality analysis supports APOBEC as a major source of mutations in head and neck squamous cell carcinoma. Oral oncology 74, 8–14.

Gillison, M.L., Akagi, K., Xiao, W., Jiang, B., Pickard, R.K.L., Li, J., Swanson, B.J., Agrawal, A.D., Zucker, M., Stache-Crain, B., et al. (2019). Human papillomavirus and the landscape of secondary genetic alterations in oral cancers. Genome Res 29, 1–17.

Glaser, A.P., Fantini, D., Wang, Y., Yu, Y., Rimar, K.J., Podojil, J.R., Miller, S.D., and Meeks, J.J. (2018). APOBEC-mediated mutagenesis in urothelial carcinoma is associated with improved survival, mutations in DNA damage response genes, and immune response. Oncotarget 9, 4537–4548.

Grompe, M., Overturf, K., al-Dhalimy, M., and Finegold, M. (1998). Therapeutic trials in the murine model of hereditary tyrosinaemia type I: a progress report. J Inherit Metab Dis 21, 518–531.

Hanahan, D., and Weinberg, R.A. (2011). Hallmarks of cancer: the next generation. Cell 144, 646–674.

Harris, R.S., and Dudley, J.P. (2015). APOBECs and virus restriction. Virology 479-480C, 131–145.

Helleday, T., Eshtad, S., and Nik-Zainal, S. (2014). Mechanisms underlying mutational signatures in human cancers. Nat Rev Genet 15, 585–598.

Henderson, S., Chakravarthy, A., Su, X., Boshoff, C., and Fenton, T.R. (2014). APOBEC-mediated cytosine deamination links PIK3CA helical domain mutations to human papillomavirus-driven tumor development. Cell Rep 7, 1833–1841.

Hultquist, J.F., Lengyel, J.A., Refsland, E.W., LaRue, R.S., Lackey, L., Brown, W.L., and Harris, R.S. (2011). Human and rhesus APOBEC3D, APOBEC3F, APOBEC3G, and APOBEC3H demonstrate a conserved capacity to restrict Vif-deficient HIV-1. J Virol 85, 11220–11234.

Ito, F., Fu, Y., Kao, S.A., Yang, H., and Chen, X.S. (2017). Family-wide comparative analysis of cytidine and methylcytidine deamination by eleven human APOBEC proteins. J Mol Biol 429, 1787–1799.

Jarvis, M.C., Ebrahimi, D., Temiz, N.A., and Harris, R.S. (2018). Mutation signatures including APOBEC in cancer cell lines. JNCI Cancer Spectr 2.

Keng, V.W., Sia, D., Sarver, A.L., Tschida, B.R., Fan, D., Alsinet, C., Sole, M., Lee, W.L., Kuka, T.P., Moriarity, B.S., et al. (2013). Sex bias occurrence of hepatocellular carcinoma in Poly7 molecular subclass is associated with EGFR. Hepatology 57, 120–130.

Keng, V.W., Tschida, B.R., Bell, J.B., and Largaespada, D.A. (2011). Modeling hepatitis B virus X-induced hepatocellular carcinoma in mice with the Sleeping Beauty transposon system. Hepatology 53, 781–790.

Keng, V.W., Villanueva, A., Chiang, D.Y., Dupuy, A.J., Ryan, B.J., Matise, I., Silverstein, K.A., Sarver, A., Starr, T.K., Akagi, K., et al. (2009). A conditional transposon-based insertional mutagenesis screen for genes associated with mouse hepatocellular carcinoma. Nat Biotechnol 27, 264–274.

Koning, F.A., Goujon, C., Bauby, H., and Malim, M.H. (2011). Target cell-mediated editing of HIV-1 cDNA by APOBEC3 proteins in human macrophages. J Virol 85, 13448–13452.

Kouno, T., Silvas, T.V., Hilbert, B.J., Shandilya, S.M.D., Bohn, M.F., Kelch, B.A., Royer, W.E., Somasundaran, M., Kurt Yilmaz, N., Matsuo, H., et al. (2017). Crystal structure of APOBEC3A bound to single-stranded DNA reveals structural basis for cytidine deamination and specificity. Nat Commun 8, 15024.

Land, A.M., Law, E.K., Carpenter, M.A., Lackey, L., Brown, W.L., and Harris, R.S. (2013). Endogenous APOBEC3A DNA cytosine deaminase is cytoplasmic and nongenotoxic. J Biol Chem 288, 17253–17260.

Landry, S., Narvaiza, I., Linfesty, D.C., and Weitzman, M.D. (2011). APOBEC3A can activate the DNA damage response and cause cell-cycle arrest. EMBO Rep 12, 444–450.

Law, E.K., Sieuwerts, A.M., LaPara, K., Leonard, B., Starrett, G.J., Molan, A.M., Temiz, N.A., Vogel, R.I., Meijer-van Gelder, M.E., Sweep, F.C., et al. (2016). The DNA cytosine deaminase APOBEC3B promotes tamoxifen resistance in ER-positive breast cancer. Sci Adv 2, e1601737.

Lawrence, C.W. (2002). Cellular roles of DNA polymerase zeta and Rev1 protein. DNA Repair (Amst) 1, 425–435.

Lefebvre, C., Bachelot, T., Filleron, T., Pedrero, M., Campone, M., Soria, J.C., Massard, C., Levy, C., Arnedos, M., Lacroix-Triki, M., et al. (2016). Mutational profile of metastatic breast cancers: a retrospective analysis. PLoS Med 13, e1002201.

Leonard, B., McCann, J.L., Starrett, G.J., Kosyakovsky, L., Luengas, E.M., Molan, A.M., Burns, M.B., McDougle, R.M., Parker, P.J., Brown, W.L., et al. (2015). The PKC/NF-kappaB signaling pathway induces APOBEC3B expression in multiple human cancers. Cancer Res 75, 4538–4547.

Lucifora, J., Xia, Y., Reisinger, F., Zhang, K., Stadler, D., Cheng, X., Sprinzl, M.F., Koppensteiner, H., Makowska, Z., Volz, T., et al. (2014). Specific and nonhepatotoxic degradation of nuclear hepatitis B virus cccDNA. Science 343, 1221–1228.

MacMillan, A.L., Kohli, R.M., and Ross, S.R. (2013). APOBEC3 inhibition of mouse mammary tumor virus infection: the role of cytidine deamination versus inhibition of reverse transcription. J Virol 87, 4808–4817.

Malim, M.H., and Bieniasz, P.D. (2012). HIV restriction factors and mechanisms of evasion. Cold Spring Harbor perspectives in medicine 2, a006940.

Marusawa, H., and Chiba, T. (2010). Helicobacter pylori-induced activation-induced cytidine deaminase expression and carcinogenesis. Curr Opin Immunol 22, 442–447.

Maruyama, W., Shirakawa, K., Matsui, H., Matsumoto, T., Yamazaki, H., Sarca, A.D., Kazuma, Y., Kobayashi, M., Shindo, K., and Takaori-Kondo, A. (2016). Classical NF-kappaB pathway is responsible for APOBEC3B expression in cancer cells. Biochem Biophys Res Commun 478, 1466–1471.

Moolenbeek, C., and Ruitenberg, E.J. (1981). The “Swiss roll”: a simple technique for histological studies of the rodent intestine. Lab Anim 15, 57–59.

Mori, S., Takeuchi, T., Ishii, Y., and Kukimoto, I. (2015). Identification of APOBEC3B promoter elements responsible for activation by human papillomavirus type 16 E6. Biochem Biophys Res Commun 460, 555–560.

Mori, S., Takeuchi, T., Ishii, Y., Yugawa, T., Kiyono, T., Nishina, H., and Kukimoto, I. (2017). Human Papillomavirus 16 E6 upregulates APOBEC3B via the TEAD transcription factor. J Virol 91.

Moser, A.R., Pitot, H.C., and Dove, W.F. (1990). A dominant mutation that predisposes to multiple intestinal neoplasia in the mouse. Science 247, 322–324.

Munteanu, I., and Mastalier, B. (2014). Genetics of colorectal cancer. J Med Life 7, 507–511.

Nik-Zainal, S., Alexandrov, L.B., Wedge, D.C., Van Loo, P., Greenman, C.D., Raine, K., Jones, C., Hinton, J., Marshall, J., Stebbings, L.A., et al. (2012). Mutational processes molding the genomes of 21 breast cancers. Cell 149, 979–993.

Nik-Zainal, S., Davies, H., Staaf, J., Ramakrishna, M., Glodzik, D., Zou, X., Martincorena, I., Alexandrov, L.B., Martin, S., Wedge, D.C., et al. (2016). Landscape of somatic mutations in 560 breast cancer whole-genome sequences. Nature 534, 47–54.

Okazaki, I.M., Hiai, H., Kakazu, N., Yamada, S., Muramatsu, M., Kinoshita, K., and Honjo, T. (2003). Constitutive expression of AID leads to tumorigenesis. J Exp Med 197, 1173–1181.

Olson, M.E., Harris, R.S., and Harki, D.A. (2018). APOBEC enzymes as targets for virus and cancer therapy. Cell chemical biology 25, 36–49.

Refsland, E.W., Stenglein, M.D., Shindo, K., Albin, J.S., Brown, W.L., and Harris, R.S. (2010). Quantitative profiling of the full *APOBEC3* mRNA repertoire in lymphocytes and tissues: implications for HIV-1 restriction. Nucleic Acids Res 38, 4274–4284.

Robbiani, D.F., and Nussenzweig, M.C. (2013). Chromosome translocation, B cell lymphoma, and activation-induced cytidine deaminase. Annu Rev Pathol 8, 79–103.

Roberts, S.A., and Gordenin, D.A. (2014). Clustered and genome-wide transient mutagenesis in human cancers: Hypermutation without permanent mutators or loss of fitness. Bioessays.

Roberts, S.A., Lawrence, M.S., Klimczak, L.J., Grimm, S.A., Fargo, D., Stojanov, P., Kiezun, A., Kryukov, G.V., Carter, S.L., Saksena, G., et al. (2013). An APOBEC cytidine deaminase mutagenesis pattern is widespread in human cancers. Nat Genet 45, 970–976.

Sasatani, M., Xi, Y., Kajimura, J., Kawamura, T., Piao, J., Masuda, Y., Honda, H., Kubo, K., Mikamoto, T., Watanabe, H., et al. (2017). Overexpression of Rev1 promotes the development of carcinogen-induced intestinal adenomas via accumulation of point mutation and suppression of apoptosis proportionally to the Rev1 expression level. Carcinogenesis 38, 570–578.

Shaban, N.M., Shi, K., Lauer, K.V., Carpenter, M.A., Richards, C.M., Salamango, D., Wang, J., Lopresti, M.W., Banerjee, S., Levin-Klein, R., et al. (2018). The antiviral and cancer genomic DNA deaminase APOBEC3H is regulated by an RNA-mediated dimerization mechanism. Mol Cell 69, 75–86 e79.

Shi, K., Carpenter, M.A., Banerjee, S., Shaban, N.M., Kurahashi, K., Salamango, D.J., McCann, J.L., Starrett, G.J., Duffy, J.V., Demir, O., et al. (2017). Structural basis for targeted DNA cytosine deamination and mutagenesis by APOBEC3A and APOBEC3B. Nat Struct Mol Biol 24, 131–139.

Sieuwerts, A.M., Schrijver, W.A., Dalm, S.U., de Weerd, V., Moelans, C.B., Ter Hoeve, N., van Diest, P.J., Martens, J.W., and van Deurzen, C.H. (2017). Progressive APOBEC3B mRNA expression in distant breast cancer metastases. PloS one 12, e0171343.

Sieuwerts, A.M., Willis, S., Burns, M.B., Look, M.P., Meijer-Van Gelder, M.E., Schlicker, A., Heideman, M.R., Jacobs, H., Wessels, L., Leyland-Jones, B., et al. (2014). Elevated APOBEC3B correlates with poor outcomes for estrogen-receptor-positive breast cancers. Hormones & cancer 5, 405–413.

Simon, V., Bloch, N., and Landau, N.R. (2015). Intrinsic host restrictions to HIV-1 and mechanisms of viral escape. Nat Immunol 16, 546–553.

Siriwardena, S.U., Chen, K., and Bhagwat, A.S. (2016). Functions and malfunctions of mammalian DNA-cytosine deaminases. Chem Rev 116, 12688–12710.

Siriwardena, S.U., Perera, M.L.W., Senevirathne, V., Stewart, J., and Bhagwat, A.S. (2019). A tumor-promoting phorbol ester causes a large increase in APOBEC3A expression and a moderate increase in APOBEC3B expression in a normal human keratinocyte cell line without increasing genomic uracils. Mol Cell Biol 39.

Stavrou, S., Crawford, D., Blouch, K., Browne, E.P., Kohli, R.M., and Ross, S.R. (2014). Different modes of retrovirus restriction by human APOBEC3A and APOBEC3G in vivo. PLoS Pathog 10, e1004145.

Stenglein, M.D., Burns, M.B., Li, M., Lengyel, J., and Harris, R.S. (2010). APOBEC3 proteins mediate the clearance of foreign DNA from human cells. Nat Struct Mol Biol 17, 222–229.

Swanton, C., McGranahan, N., Starrett, G.J., and Harris, R.S. (2015). APOBEC enzymes: mutagenic fuel for cancer evolution and heterogeneity. Cancer Discov 5, 704–712.

Thielen, B.K., McNevin, J.P., McElrath, M.J., Hunt, B.V., Klein, K.C., and Lingappa, J.R. (2010). Innate immune signaling induces high levels of TC-specific deaminase activity in primary monocyte-derived cells through expression of APOBEC3A isoforms. J Biol Chem 285, 27753–27766.

Venkatesan, S., Rosenthal, R., Kanu, N., McGranahan, N., Bartek, J., Quezada, S.A., Hare, J., Harris, R.S., and Swanton, C. (2018). Perspective: APOBEC mutagenesis in drug resistance and immune escape in HIV and cancer evolution. Annals of oncology : official journal of the European Society for Medical Oncology / ESMO 29, 563–572.

Vieira, V.C., Leonard, B., White, E.A., Starrett, G.J., Temiz, N.A., Lorenz, L.D., Lee, D., Soares, M.A., Lambert, P.F., Howley, P.M., et al. (2014). Human papillomavirus E6 triggers upregulation of the antiviral and cancer genomic DNA deaminase APOBEC3B. MBio 5.

Walker, B.A., Wardell, C.P., Murison, A., Boyle, E.M., Begum, D.B., Dahir, N.M., Proszek, P.Z., Melchor, L., Pawlyn, C., Kaiser, M.F., et al. (2015). APOBEC family mutational signatures are associated with poor prognosis translocations in multiple myeloma. Nat Commun 6, 6997.

Wangensteen, K.J., Wilber, A., Keng, V.W., He, Z., Matise, I., Wangensteen, L., Carson, C.M., Chen, Y., Steer, C.J., McIvor, R.S., et al. (2008). A facile method for somatic, lifelong manipulation of multiple genes in the mouse liver. Hepatology 47, 1714–1724.

Warren, C.J., Xu, T., Guo, K., Griffin, L.M., Westrich, J.A., Lee, D., Lambert, P.F., Santiago, M.L., and Pyeon, D. (2015). APOBEC3A functions as a restriction factor of human papillomavirus. J Virol 89, 688–702.

Wilber, A., Frandsen, J.L., Wangensteen, K.J., Ekker, S.C., Wang, X., and McIvor, R.S. (2005). Dynamic gene expression after systemic delivery of plasmid DNA as determined by in vivo bioluminescence imaging. Hum Gene Ther 16, 1325–1332.

Xu, L., Chang, Y., An, H., Zhu, Y., Yang, Y., and Xu, J. (2015). High APOBEC3B expression is a predictor of recurrence in patients with low-risk clear cell renal cell carcinoma. Urologic oncology 33, 340 e341–348.

Yamanaka, S., Balestra, M.E., Ferrell, L.D., Fan, J., Arnold, K.S., Taylor, S., Taylor, J.M., and Innerarity, T.L. (1995). Apolipoprotein B mRNA-editing protein induces hepatocellular carcinoma and dysplasia in transgenic animals. Proc Natl Acad Sci U S A 92, 8483–8487.

Yan, S., He, F., Gao, B., Wu, H., Li, M., Huang, L., Liang, J., Wu, Q., and Li, Y. (2016). Increased APOBEC3B predicts worse outcomes in lung cancer: a comprehensive retrospective study. Journal of Cancer 7, 618–625.

Zor, T., and Selinger, Z. (1996). Linearization of the Bradford protein assay increases its sensitivity: theoretical and experimental studies. Analytical biochemistry 236, 302–308.

