## Supplementary Methods and Figures S1-S12 for "APOBEC3A Catalyzes Mutation and Drives Carcinogenesis *In Vivo*"

### SUPPLEMENTAL INFORMATION

#### METHODS

**Animal care.** Mice were housed at the University of Minnesota Twin Cities, University of Pennsylvania, and University of Illinois at Chicago animal facilities under specific pathogen free conditions. All studies were performed in accordance with the recommendations in the Guide for the Care and Use of Laboratory Animals of the National Institutes of Health. The experiments performed with mice in this study were approved by the University of Pennsylvania Institutional Animal Care and Use Committee (protocol #805690), University of Illinois at Chicago Animal Care Committee (protocol #18–168), and University of Minnesota Institutional Animal Care and Use Committee (protocols 1901-36652A, 1602-33523A, 1808A36277).

**A3A Apc$^{Min}$ experiments.** A3A$^{low}$ transgenic mice on an *APOBEC3*-null background have been described (Stavrou et al., 2014). A3A$^{low}$ transgenic females were bred with Apc$^{Min}$ males (Stock 002020; Jackson Laboratory, Bar Harbor, Maine). Mice resulting from this cross were euthanized by $CO_2$ administration at the age of 4 months, and intestinal tissue was analyzed for polyp formation without knowledge of genotype information (*i.e.*, blinded). The Apc$^{Min}$ and A3A/Apc$^{Min}$ and mice shown in Figure S5 were additionally heterozygous for a deletion in mouse *APOBEC3*.

***A3B* transgenesis.** pTraffic was modified by replacing IRES-GFP with IRES-Firefly-Luciferase using *Xho*I and *Bsu*36I cut sites. A3Bi (A3B cDNA containing L1 intron) was then amplified from pCDNA3.1-A3Bi (Hultquist et al., 2011) using primers A3B_*Asc*I_F 5' – NNNNGGCGCGCCACCATGAATCCACAGATC – 3' and A3B_*Nhe*I_R 5' – NNNNGCTAGCTCAGTTTCCCTGATTCTGGAGAATGG – 3', digested with *Asc*I and *Nhe*I and inserted into pTraffic-IRES-FLuciferase. Cre dependent expression of A3B was verified by immunoblot of HEK-293T cells transfected with pTraffic-IRES-FLuciferase-*Asc*I-A3Bi-*Nhe*I with or without cotransfection of a Cre expressing plasmid [antibodies – β-Actin (CST #4970), Luciferase (Novus NB600-307), DsRed (Clontech 632392), and A3B (Brown et al., 2019)]. A3B-DsRed transgenic mice were prepared by pronuclear injection of pTraffic-IRES-FLuciferase-AscI-A3Bi-NheI plasmid into a C57Bl/6 background. DsRed fluorescence was visualized in tail snips on a Leica DMIL microscope. A3B-NTD, A3B-CTD, and DsRed DNA fragments were amplified with the following primer sets: NTD_F 5' – CCACAGATCAGAAATCCGATGGA – 3', NTD_R 5' – CTCGAGACTAAAGGCAACAGTGCTG – 3'; CTD_F 5' – GCCAGTGACTAGTGCTGCAAG – 3', CTD_R 5' – GTTTCCCTGATTCTGGAGAATGG – 3'; DsRed_F 5' – CCCATGGTCTTCTTCTGCAT – 3', DsRed_R 5' – AAGGTGTACGTGAAGCACCC – 3'. A3B-NTD primers were used for genotyping. The DsRed PCR fragment was cloned using CloneJET PCR Cloning Kit (Thermo Fisher Scientific) and Sanger sequenced.

**A3 *Fah* experiments.** *Fah*-deficient mice expressing SB11 (*Fah*$^{-/-}$; *Rosa26-SB11*$^{Tg/WT}$) were generated and maintained with drinking water containing 7.5 µg/mL NTBC (Sigma Aldrich) as described (Keng et al., 2011) until introduction of transposon vectors by hydrodynamic tail vein injections at 8-10 weeks of age. Intron-containing human A3 cDNAs were ordered as gBlocks (IDT) and cloned into the pENTR entry vector (Invitrogen) using *Not*I-*Nco*I (A3Ai, A3Bi, A3Ci, A3Gi), *Mfe*I-*Not*I (A3Di, A3Fi), or *Kpn*I-*Not*I (A3Hi). The final Gateway destination plasmid co-expressing *Fah*, GFP, and Luciferase (Keng et al., 2013) was combined with pENTR-A3 vectors using Gateway LR clonase mix (#11791-020; Thermo Fisher Scientific) to generate pT2/GD-A3 delivery plasmids. The transposon vector expressing a shRNA against *Tp53* and its validation *in vivo* has been described (Wangensteen et al., 2008).

Immediately prior to hydrodynamic deliver, 8-10 week-old *Fah*-null SB11 mice were anesthetized by administrating 25 μL anesthetic cocktail (8 $mg/mL^{-1}$ ketamine HCl, 0.1 $mg/mL^{-1}$ acepromazine maleate and 0.01 $mg/mL^{-1}$ butorphanol tartrate) i.p. Each animal was injected with 20 μg of each transposon plasmid (PureLink™ HiPure Plasmid Filter Maxiprep kit, Invitrogen) diluted in lactated Ringer's solution (Fisher Scientific) to an injection volume based on the weight of the mouse (10% vol/wt). Immediately following injection, animals were placed on normal drinking water to promote liver repopulation with *Fah-A3* transgenic cells. Luciferase expression in repopulated transgenic livers was monitored using the Xenogen IVIS 100 (Perkin Elmer) as described (Wilber et al., 2005). Briefly, animals were injected with 93-150 mg luciferin/kg body weight (Xenogen) i.p. 10-15 minutes prior to imaging. Mice were anesthetized using 1-1.5% isoflurane and $O_2/N_2O$ via nose cone, positioned in a custom-built cradle, and imaged for 1-5 minutes.

6 months post-injection, mice were euthanized and weighed and liver tissues were harvested and weighed. Liver mass was recorded and all visible distinct nodules (>2 mm in diameter) were counted and carefully isolated from neighboring tumor-free tissue. All nodules >2 mm from were halved for DNA and RNA isolation. Genomic DNA isolations were performed using either AllPrep DNA/RNA kit or DNeasy Blood & Tissue kit per manufacturer protocols (QIAGEN). RNA isolations were conducted with either the AllPrep DNA/RNA kit or RNeasy Mini Kit following manufacture protocols (QIAGEN).

Protein lysates were obtained through vortex mediated homogenization with stainless steel beads (QIAGEN) for two minutes at 4°C in protein lysis buffer (50 mM Tris-Cl pH 7.4, 250 mM NaCl, 0.5% Igepal NP-40) at 10-20x vol/wt tissue. Homogenized lysates were cleared of debris through centrifugation (max speed for 20 min at 4°C) and quantified with a NanoDrop (ThermoFisher). Samples were treated with RNaseA (1.67 μg/μL) for 10 minutes at room temperature, then combined 1:1 with SDS-PAGE loading buffer (62.5 mM Tris-Cl pH6.8, 20% glycerol, 7.5% SDS, 5% 2-mercaptoethanol, and 250 mM DTT). Proteins were separated by a 12.5% SDS-PAGE gel and transferred to PVDF-FL membranes (Millipore). Membranes were blocked in blocking solution (5% milk + PBS supplemented with 0.1% Tween20) and then incubated with primary antibody diluted in blocking solution. Primary immunoblotting antibodies were mouse α-Tubulin (Sigma Aldrich T5168), rabbit α-human A3A/B/G mAb [mAb 5210-87-13 (Brown et al., 2019)], rabbit α-human A3C pAb (Proteintech 10591-1-AP), rabbit α-human A3H pAb (Novus NBP1-91682), and rabbit α-firefly luciferase pAb (Abcam ab21176). Secondary antibodies were diluted in blocking solution supplemented with 0.02% SDS. Secondary antibodies used for detection were α-rabbit 800CW (LI-COR 827-08365), α-mouse 680LT (LI-COR 925-68020), α-rabbit HRP (Cell Signaling 7074P2), and α-mouse HRP (Cell Signaling 7076P2). Membranes were imaged with an Odyssey Classic scanner and Odyssey Fc imager (LI-COR).

**A3A$^{high}$ transgene characterization.** RNA from the intestines of WT, A3A$^{high}$, and A3A$^{low}$ animals was extracted and RNA-seq libraries were prepared and sequenced on a HiSeq 2500 125x2 bp. Reads were mapped to the mouse genome with human A3A cDNA appended as an additional chromosome using tophat. Read coverage plots suggested a 3' truncation.

For further mapping of the 3' truncation, spleens were harvested from 3-month-old mice. Splenic DNA was isolated using the DNeasy Blood and Tissue Kit (Qiagen). RNA was isolated with the use of Trizol (Invitrogen), and was further processed according to the Qiagen RNA cleanup protocol (treated with DNaseI to eliminate any contaminating genomic DNA). Purified RNA was converted to cDNA using the SuperScript III First Strand Synthesis System for RT-PCR using 50uM oligo(dT)20 as the primer (Invitrogen). Full-length cDNA was obtained by rapid amplification of cDNA ends (RACE) using *A3A* transgene forward primer 5'-TGGACCTGGTTCCTTCTTT, the poly(A) tail functioning as the 3'end tag. Bands were excised and the fragments were cloned into pCR2.1-TOPO vector as specified by the manufacturer (Invitrogen). Additionally, the transgene DNA was amplified using primers CAG promoter 5'-GGGCGGGGTTCGGCTTCTGGCGTGTGAC; and CAG polyA tail 5'-CAGGGCATTGGCCACACCAGCCACCACC. Both the cloned cDNA and amplified genomic DNA were Sanger sequenced.

**RT-PCR and qPCR.** RNA was extracted from mouse tissues (Fah and $Apc^{Min}$) with RNeasy mini kit (QIAGEN) and from polyps with Allprep DNA/RNA mini kit (QIAGEN). cDNA was prepared using Transcriptor reverse transcriptase (Roche) with random hexamer priming. Full length *A3A* transcript was amplified with the following primers – A3A_full_length-F 5' – ATGGAAGCCAGCCCAGCATC; A3A_full_length-R 5' – GTTTCCCTGATTCTGGAGAATGG. Relative transcript levels were measured by qPCR with LightCycler® 480 Probes Master mix (Roche) on a LightCycler® 480 instrument (Roche) and the following primers: *A3A* [in conjunction with UPL probe 97 (Roche)] A3A_qPCR_F 5' - CCACACATATTCACTTCCAACT, A3A_qPCR_R 5' – TGTGCTGGTCCATCTTGA; *Tbp* [in conjunction with UPL probe 97(Roche)] mouse_Tbp_F 5' – GGGGAGCTGTGATGTGAAGT, mouse_Tbp_R 5' – CCAGGAAATAATTCTGGCTCA. APOBEC3 mRNA was quantified with previously described primers (*A3B*, *A3D*, *A3F*, *A3G*, *TBP*) or *A3A* (Forward 5' – CGGTCAAGATGGACCAGCAC; Reverse 5' – GAAGGAACGCACGCACTTC), *A3C* (Forward 5' – AGCCAACGATCGGAACGAAA; Reverse 5' – AGGGCTCCAAGATGTGTACC), *A3H* (Forward 5' – TCAGAAGGCCTTACTACCCG; Reverse 5' – ATGAAGTCAACCAGCTCCCAG) using SsoFast EvaGreen Supermix (BioRad) per manufactures protocols. *Luciferase* mRNA was quantified using LightCycler® 480 Probes Master Mix (Roche) with primers Forward 5' - TCCATCTTGCTCCAACACCC and Reverse 5' – TCGTCTTTCCGTGCTCCAAA. All mRNA quantification was obtained using a LightCycler® 480 instrument (Roche).

**Deaminase activity assays.** Spleens from 4-month old $A3A^{low}$ mice were homogenized and lysed in HED buffer [25mM HEPES, 5mM EDTA, 10% Glycerol, 1mM DTT, 1x Protease inhibitor (cOmplete-Roche)]. Lysates were sonicated for 20 minutes in a water bath sonicator, cleared by centrifugation and concentrated using Amicon Ultra 0.5 mL centrifugal filters with a MWCO of 10k with two washes of 400 µL of buffer (25 mM HEPES pH 7.4, 15 mM EDTA, cOmplete EDTA-free protease inhibitors, 10% glycerol). Protein concentration in lysates was quantified by Bradford assay (Zor and Selinger, 1996). Oligo NUP93 (6-FAM)-GCAAGCTGTTCAGCTTGCTGA (Buisson et al., 2019) was allowed to form a hairpin by heating a 10 µM stock in 1 mM Tris-Cl and 0.1 mM EDTA to 65°C for 5 minutes then allowing the DNA to cool to room temperature. 75 µg of protein lysate was incubated with 800 nM of NUP93 hairpin oligo at 37°C with 100 µg/mL RNase A for 24 hours. 0.1 units of UDG (NEB) was added to each reaction and incubated for 10 minutes at 37°C. NaOH was added to a final concentration of 100 mM and samples were heated to 95°C for 10 minutes to cause breakage of abasic sites caused by deamination and subsequent uracil removal. Samples were separated by 15% TBE-Urea PAGE to resolve product and imaged on a Typhoon FLA 7000 biomolecular imager (GE Healthcare Life Sciences). A3A-mycHis purified from 293T cells was used at 1 nM as a positive control for deamination activity (Shi et al., 2017). APOBEC3 activity was detected similarly using lysates obtained as described above.

**Handling of intestinal tissue.** Intestines from 4-month old $Apc^{Min}$ and $A3A^{low}/Apc^{Min}$ mice were removed from duodenum to colon and placed on PBS-soaked Bibulous paper. Intestines were divided into sections and then carefully sliced lengthwise, cleaned and spread with lumen side upward. Select large polyps from the distal colon were excised and flash frozen for future molecular analysis. Intestines were fixed overnight in 10% buffered formalin solution. Polyp numbers were counted (blinded to mouse genotype) using a Leica S8 APO stereo microscope (at University of Minnesota) or a Nikon SMZ1500 (at University of Pennsylvania). Statistical differences between $Apc^{Min}$ and $A3A^{low}/Apc^{Min}$ animals were determined by two-sided Wilcoxon rank sum test using Prism 6 software.

**Histology.** The small intestine and colon of $Apc^{Min}$ and $A3A^{low}/Apc^{Min}$ mice were isolated and fixed in 10% neutral formalin. The flattened segments of intestinal tissues were rolled lengthwise into "Swiss rolls" (Moolenbeek and Ruitenberg, 1981) and embedded in paraffin (FFPE). In addition, normal livers and liver tumors generated with the A3A Fah model were collected and fixed as above. Hematoxylin and eosin (H&E) staining of the FFPE specimens was performed as follows: Tissues were sectioned at 4 μm, mounted on positively charged adhesive slides and allowed to air-dry for at least 24 hours prior to staining. To deparaffinize and rehydrate the samples, slides were baked in a 60-62°C oven for 20 min, washed 3 times with xylene for 5 min each, soaked in graded alcohols (100% × 2, 95% and 80% for 3 min each), and then rinsed in running water for 5 min. The tissues were stained with hematoxylin for 5 min, rinsed in running water for 30 sec, followed by two dips in acid solution and 30-90 sec in ammonia water (bluing solution). After a 10 min water rinse, the slides were transferred in 80% ethanol for 1 min, counterstained with eosin for 1 min, dehydrated in graded alcohols and xylene, and cover-slipped using Permount mounting media. The H&E stained slides were subsequently scanned at 40× magnification and visualized using the Aperio ScanScope XT system (Leica Biosystems) as described (Brown et al., 2019).

**Immunohistochemistry.** 4-μm thick sections of FFPE intestinal and liver tissues were mounted on positively charged, adhesive slides and allowed to air-dry for at least 24 hrs. To deparaffinize and rehydrate the samples, slides were baked in a 65°C oven for 20 min, washed 3 times with CitrisolvTM (Decon Labs, #1601) or xylene for 5-min/each, soaked in graded alcohols (100% x 2, 95% and 80% for 3 min/each), and then rinsed in running water for at least 5 min. Epitope retrieval was performed using Reveal Decloaker (BioCare Medical, #RV1000M) in a steamer for 35 min, followed by a 20 min "cool-down" period. Then, slides were rinsed with running tap water for 5 min and transferred to TBST for 5 min. Endogenous peroxidase activity was quenched by placing the slides in 3% $H_2O_2$ in TBST for 10 min at RT, followed by a 5-min rinse under running water. To block non-specific binding of primary antibody, sections were covered with Rodent Block M (BioCare Medical, # RBM961) for 15 min at RT. After blocking, sections of each specimen were incubated overnight at 4°C with a rabbit α-human A3A/B/G mAb [5210-87-13 (Brown et al., 2019)] diluted 1:350 in 10% Rodent Block M in TBST.

Following overnight incubation with primary antibody, sections were rinsed in TBST for 5 min, and completely covered with anti-rabbit poly-HRP-IgG (Leica Biosystems, Novolink Polymer, #RE7260-K) for 30 min at RT. The reaction product was developed using the Novolink DAB substrate kit (Leica Biosystems, # RE7230-K) at RT for 3 min, rinsed in tap water for 5 min, counterstained in Mayer's hematoxylin solution (Electron Microscopy Sciences, # 26252-01) for 5 min, dehydrated in graded alcohols and CitrisolvTM, and cover-slipped using Permount mounting media. Nuclear and cytoplasmic A3A immunoreactivity was visualized using the Aperio ScanScope XT (Leica Biosystems).

**Exome library sequencing and analysis of mutational patterns.** Genomic DNA was prepared from frozen colon polyp tissue using Allprep DNA/RNA mini kit (QIAGEN). Matched normal DNA was extracted from tail snips following the Gentra Puregene Tissue DNA isolation protocols (Qiagen). DNA was sheared and adapters added using SureSelectQXT Library Prep kit (Agilent) and exonic DNA fragments were captured using SureSelectXT Mouse All Exon kit (Agilent). $Apc^{Min}$ libraries were sequenced 75x2 bp on a NextSeq 550 instrument (Illumina) to a target depth of 75x coverage for tumor samples and 30x coverage for matched normal. Fah libraries were sequenced 150x2 PE on a NovaSeq 6000 (Illumina) to similar read depths. Resulting sequences were aligned to the mouse genome (mm10) using the Burrows-Wheeler Aligner (version 0.7.17). PCR duplicates were removed using Picard (version 2.18.16). Reads were locally realigned around InDels using GATK3 (version 3.6.0) tools RealignerTargetCreator to create intervals, followed by IndelRealigner on the aligned bam files. The exome-seq alignments from the normal tail tissues were used to create a "panel of normals" background set using MuTect2 from GATK3 (version 3.6.0) in artifact detection mode. Mutations were called from the polyp samples compared to the matched normal, including the "panel of normals" as background using MuTect2 from GATK3. Single nucleotide variants (SNVs) that passed the internal GATK3 filter were used for downstream analysis. Mutational patterns were analyzed in R (version 3.6.0) using the "MutationalPatterns" package (Blokzijl et al., 2018).

COSMIC single base substitution mutational signatures (v3 – May 2019 https://cancer.sanger.ac.uk/cosmic/signatures/SBS/) were obtained from https://www.synapse.org/#!Synapse:syn11738319. *De-novo* non-negative matrix factorization of mutational signatures was performed with the "extract_signature" command, with a rank of 2 and 10 iterations. TCW mutation enrichment score was calculated as described (Jarvis et al., 2018). Sequence logos of -2 to +2 sequence surrounding C-to-T mutations was created using "ggseqlogo" package in R. Prediction of mutation effect on protein sequence was done with SnpEff (version 4.3t) using the SnpEff mm10 database. Loss of heterozygosity of chromosome 18 was calculated by comparing tumor to matched normal alignments using Varscan (version 2.3.9) in somatic mode.

**TCGA variant and Euclidean distance analysis.** All whole exome sequencing variant information from The Cancer Genome Atlas (TCGA) was downloaded from the Broad GDAC firehose database (https://gdac.broadinstitute.org/). Variant information from the A3A and A3B yeast models of mutagenesis were downloaded from previous published material (Chan et al., 2015). Only samples with greater than 300 C-to-T mutations were included in the Euclidean distance and sequencing logo analyses. SNVs that were used in these analyses included only C-to-T variants in a TC dinucleotide, excluding all mutations as CG motifs. All C-to-T variants from the $Apc^{Min}$, A3A yeast model, and A3B yeast model were used in this analysis. A matrix comprised of the number of mutations within a tetranucleotide across all samples within a cancer type was generated, and counts were normalized to frequency within each cancer type. This matrix was converted to a hierarchical distance matrix using the hclust function in R, which was then plotted as dendrograms. All sequences logos were again generated and visualized using "ggseqlogo" package in R. Total mutation counts per cancer type were generated by simple counting of all C-to-T mutations at TC dinucleotide, excluding all mutations as CG motifs.

#### SUPPLEMENTAL FIGURES

**Figures S1-S12.** Please see following pages.

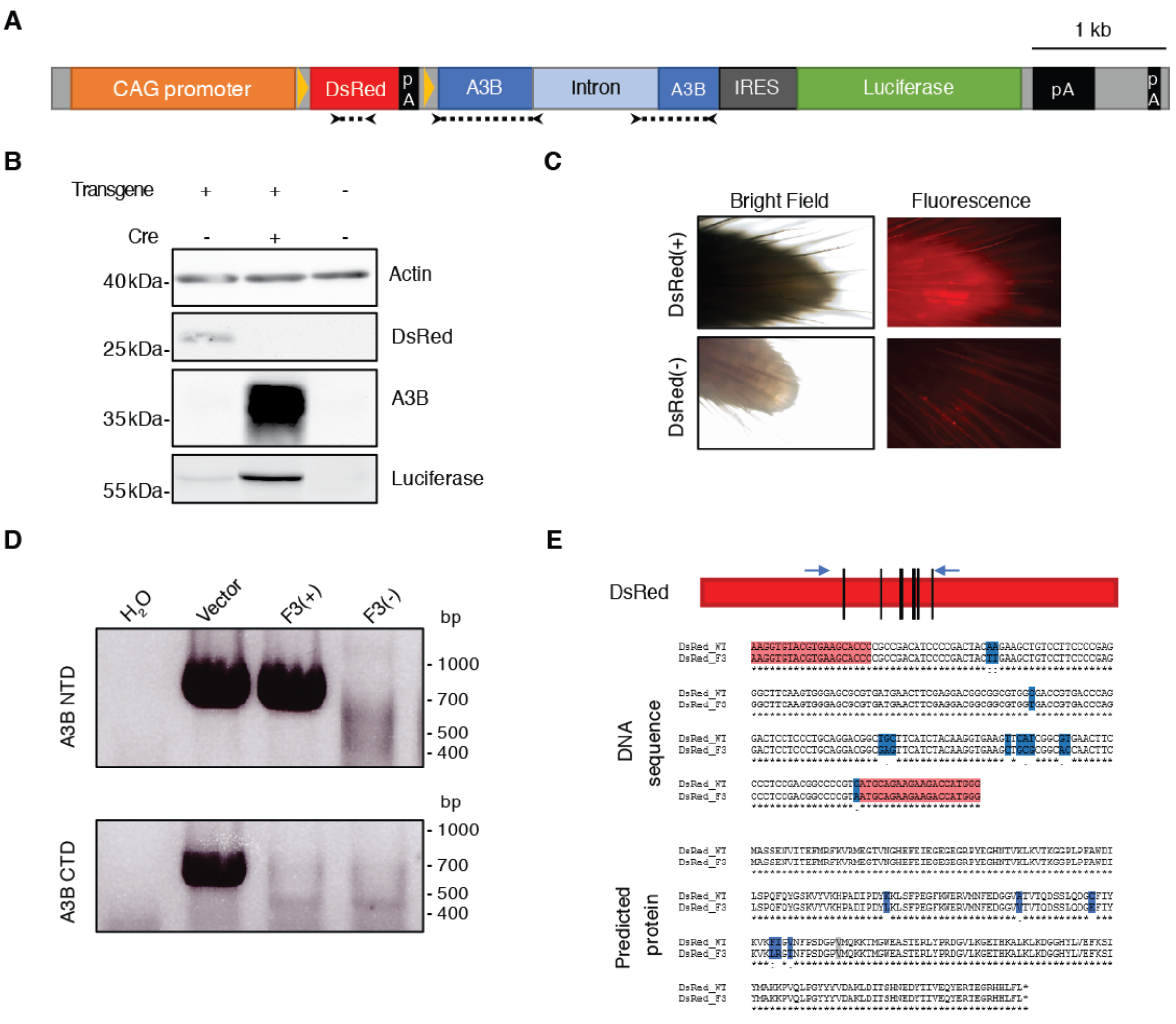


**Figure S1. Conditional A3B transgene loses activity after small number of generations.**
(**A**) Schematic of *A3B* transgene. Floxed *DsRed* gene is under the control of the CAG promoter and blocks transcription of an *A3B* minigene. Cre expression excises *DsRed* and allows expression of both A3B and firefly luciferase. Arrows and dotted lines indicate locations of primers used in panels D and E.
(**B**) Immunoblots from 293T cells either untransfected or transfected with the transgene plasmid described in panel (A), with or without the addition of a Cre expression plasmid.
(**C**) Representative images of DsRed expression in tail tips of transgenic mice compared to no fluorescence in nontransgenic mice.
(**D**)Genotyping PCR results for representative third generation animals. Mouse F3(+) was PCR-positive for the 5' end of *A3B* (encoding the N-terminal domain) and negative for the 3' end of *A3B* (encoding the C-terminal domain). This animal also failed to show fluorescence and the *DsRed* cassette was later found to harbor 13 point mutations (panel E). Mouse F3(-) was a transgene-negative littermate. Transgene vector DNA diluted in 293T genomic DNA was used as a positive control for PCR amplification.
(**E**) Mutations detected in *DsRed* fragment in F3(+) mouse described in panel (D). These 13 point mutations are predicted to lead to 6 amino acid changes (blue) and one silent change (grey).

**A**

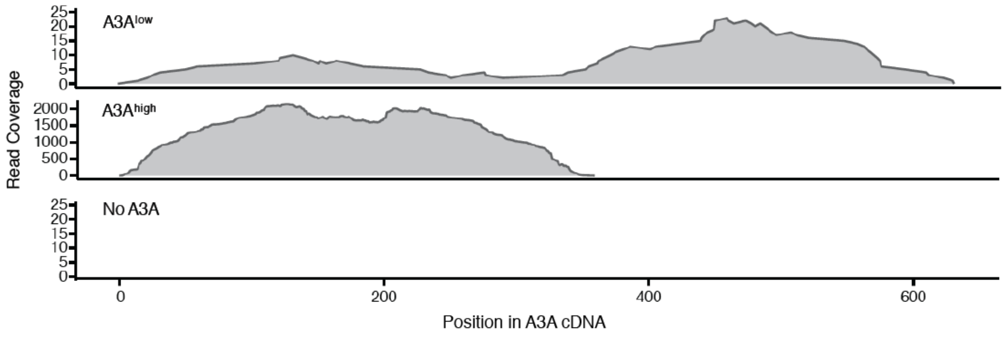


**B**

```
Native_A3A        MEASPASGPRHLMDPHIFTSNFNNGIGRHKTYLCYEVERLDNGTSVKMDQHRGFLHNQAK
A3A-high_cDNA     MEASPASGPRHLMDPHIFTSNFNNGIGRHKTYLCYEVERLDNGTSVKMDQHRGFLHNQAK
Ngly1_CTD         ------------------------------------------------------------

Native_A3A        NLLCGFYGRHAELRFLDLVPSLQLDPAQIYRVTWFISWSPCFSWGCAGEVRAFLQENTHV
A3A-high_cDNA     NLLCGFYGRHAELRFLDLVPSLQLDPAQIYRVTWFISWSPCFSWVVEVYLARKEGSSFAY
Ngly1_CTD         -------------------------------------------DWNMVYLARKEGSSFAY

Native_A3A        RLRIFAARIYDYDPLYKEALQMLRDAGAQVSIMTYDEFKHCWDTFVDHQGCPFQPWDGLD
A3A-high_cDNA     ISWKFECGSAGLKVDTVSIRTSSQSFESGSVRWKLRSETAQVNLLGDKNLRSYNDFSGAT
Ngly1_CTD         ISWKFECGSAGLKVDTVSIRTSSQSFESGSVRWKLRSETAQVNLLGDKNLRSYNDFSGAT

Native_A3A        EHSQALSGRLRAILQNQGN
A3A-high_cDNA     EVTLEAELSRGDGDVAWQHTQLFRQSLNDSGENGLEIIIITFNDL
Ngly1_CTD         EVTLEAELSRGDGDVAWQHTQLFRQSLNDSGENGLEIIIITFNDL
```

**Figure S2. A3A-A9 transgenic mice express truncated form of A3A.**
(**A**) Read coverage across *A3A* cDNA from RNA-seq libraries arising from the intestines of representative A3A$^{low}$, A3A$^{high}$, and WT mice.
(**B**) Predicted protein produced by fusion of *A3A-Ngly1* in A3A$^{high}$ transgenic animals. Amino acid prediction based upon results of 3’ RACE and Sanger sequencing.

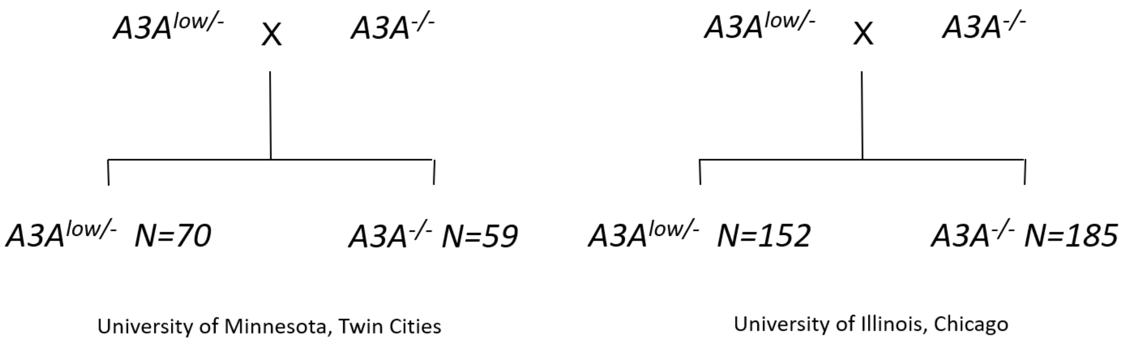


**Figure S3. Mendelian ratios for inheritance of A3A$^{low}$ transgene.**

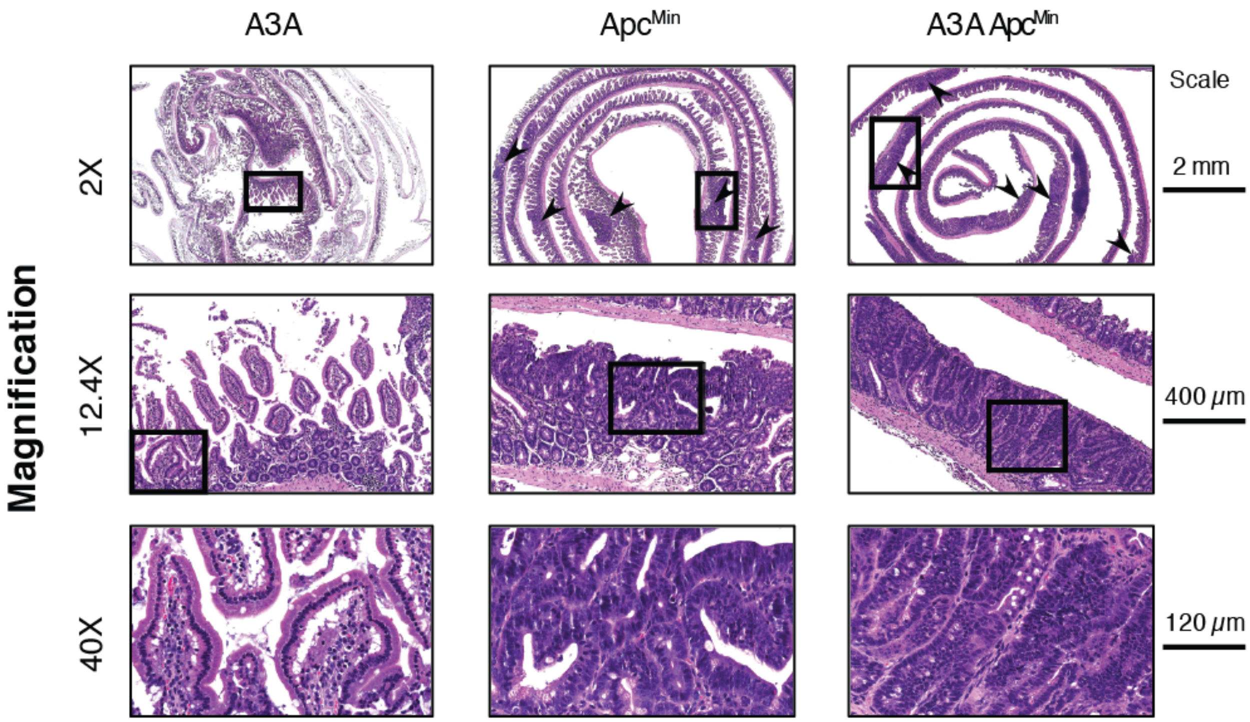


**Figure S4. Histology of small intestine section from A3A$^{low}$, Apc$^{Min}$ and A3A$^{low}$/APC$^{Min}$ mice.** Representative low- and high-power photomicrographs of indicated H&E stained small intestine sections. Black rectangle in upper low power images indicates regions shown in higher magnification in the image immediately bellow. Arrowheads in lower power magnification point to polyps.

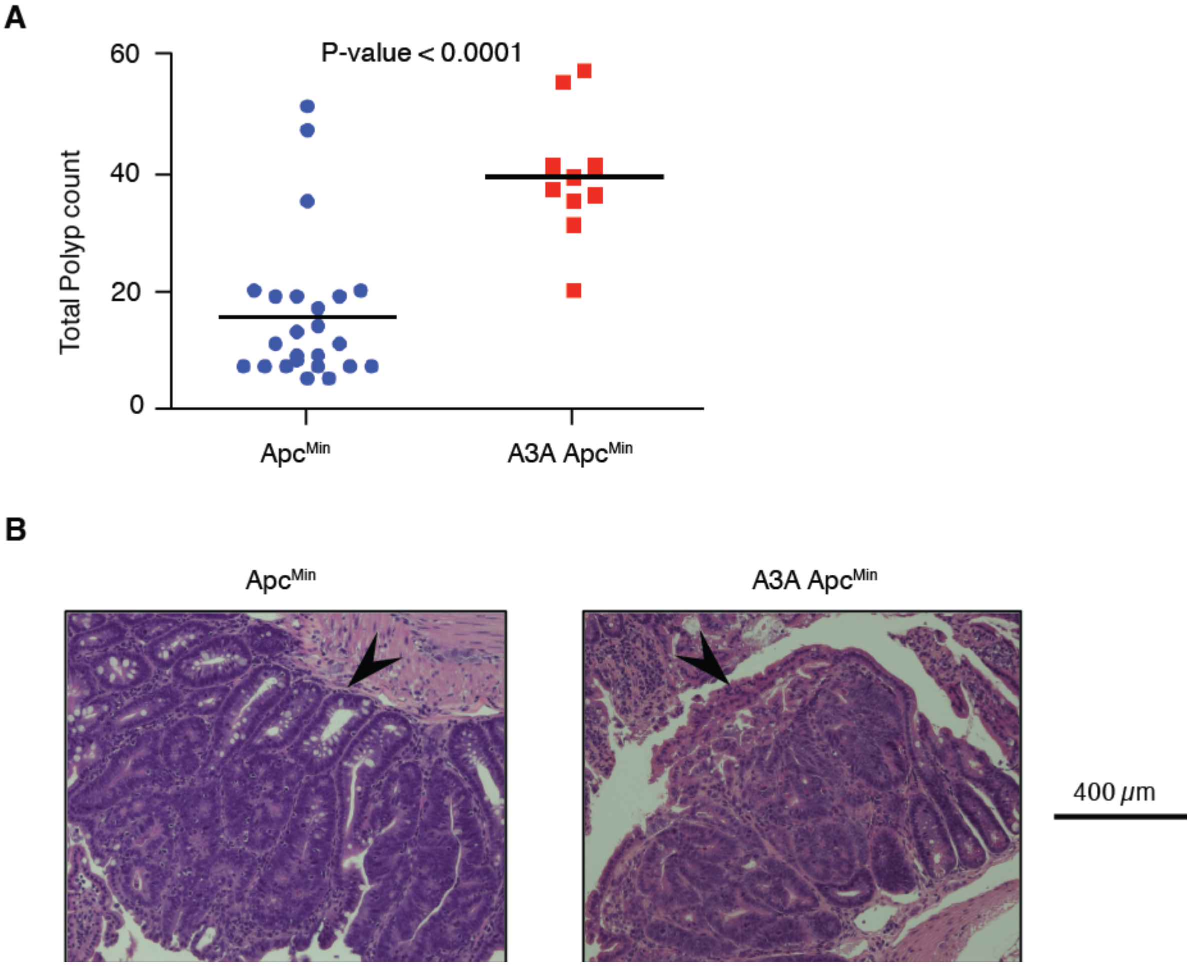


**Figure S5. A3A$^{low}$/Apc$^{Min}$ mice develop more tumors than Apc$^{Min}$ mice.**
(**A**) Numbers of intestinal polyps in mice Apc$^{Min}$ mice with or without the A3A$^{low}$ transgene. These experiments were carried out in the mouse facilities at the University of Pennsylvania, independently of the studies presented in Fig. 2.
(**B**) Representative images of H&E stained sections of small intestine from Apc$^{Min}$ and A3A$^{low}$/Apc$^{Min}$ mice from cohort in panel (A). Arrowheads point to polyps.

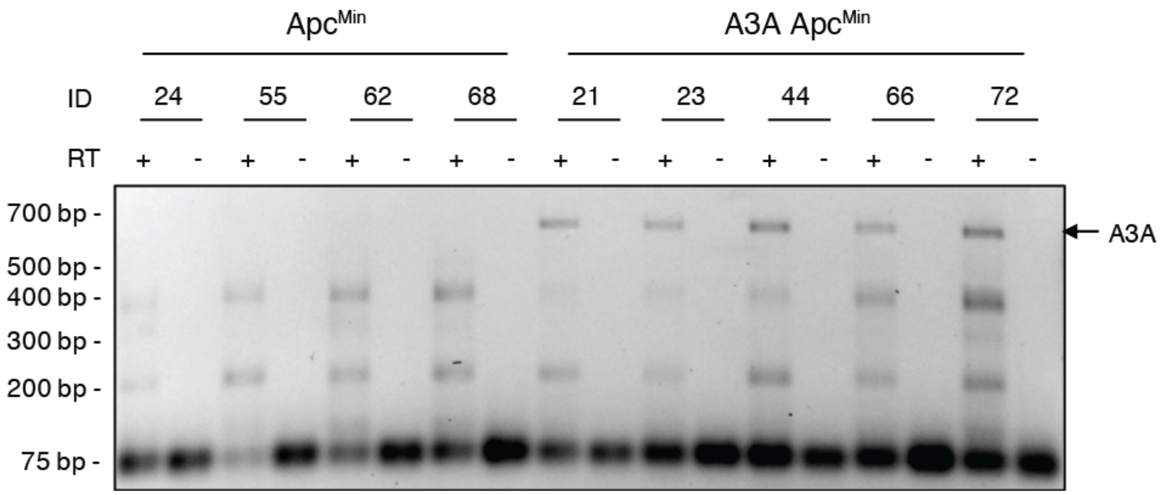


**Figure S6. RT-PCR analysis of *A3A* expression in colorectal polyps from $Apc^{Min}$ and A3A/$Apc^{Min}$ mice.**

Agarose gel image showing full-length *A3A* cDNA from the mRNA of polyps derived from $Apc^{Min}$ and A3A/$Apc^{Min}$ mice. The non-specific bands indicate similar levels of nucleic acid in these RT-PCR reactions. Parallel reactions with no RT show no genomic DNA contamination.

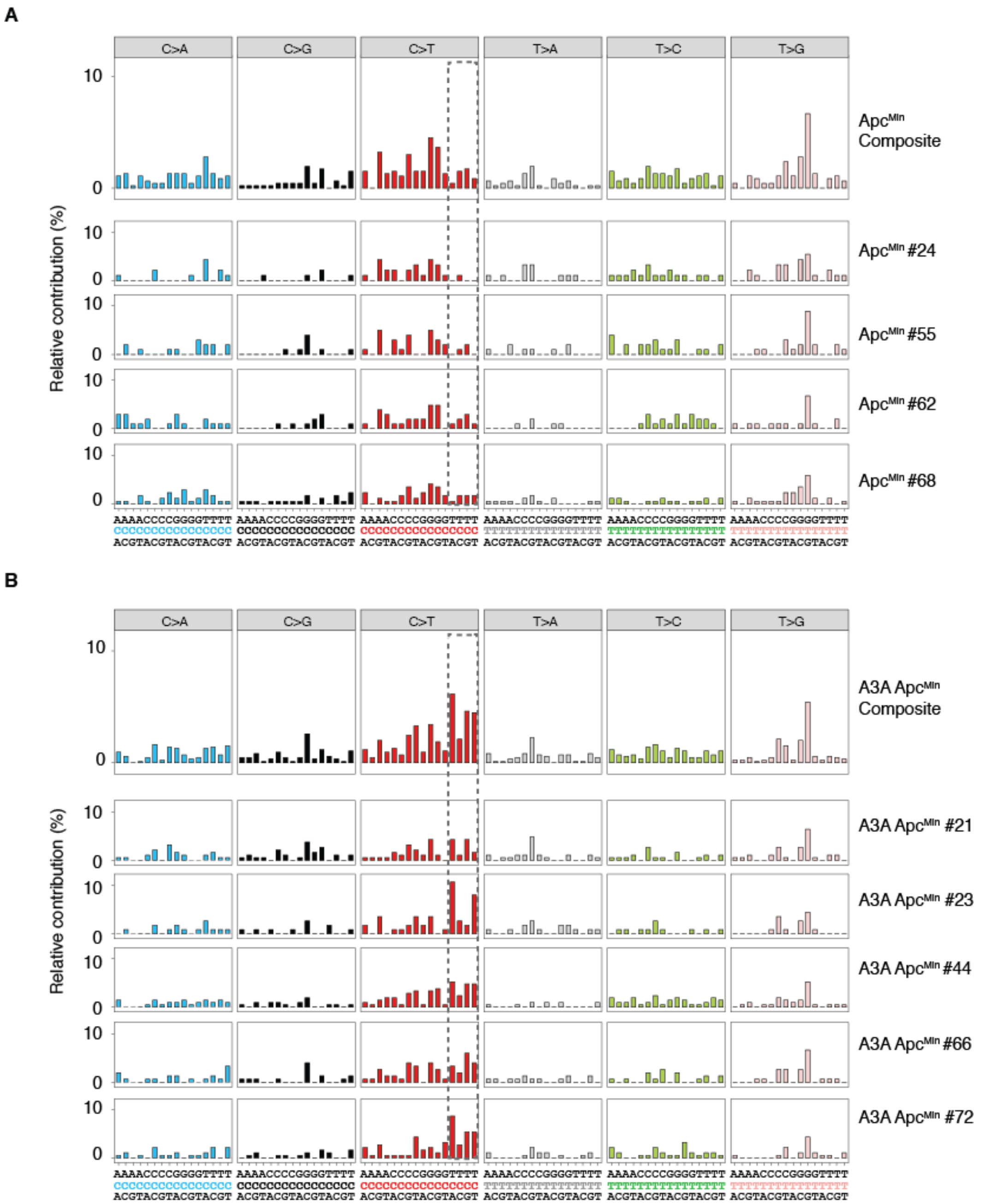


**Figure S7. Individual and composite trinucleotide mutational profiles in Apc$^{Min}$ and A3A/Apc$^{Min}$ tumors.**

(**A-B**) Barplots representing composite trinucleotide mutation profiles for all base substitutions in Apc$^{Min}$ versus A3A$^{low}$/Apc$^{Min}$ tumors, respectively (identical to Fig. 3D). Barplots for individual tumor trinucleotide mutation profiles are shown below each composite plot. Dotted box highlights the trinucleotide motifs that define the APOBEC mutation signature.

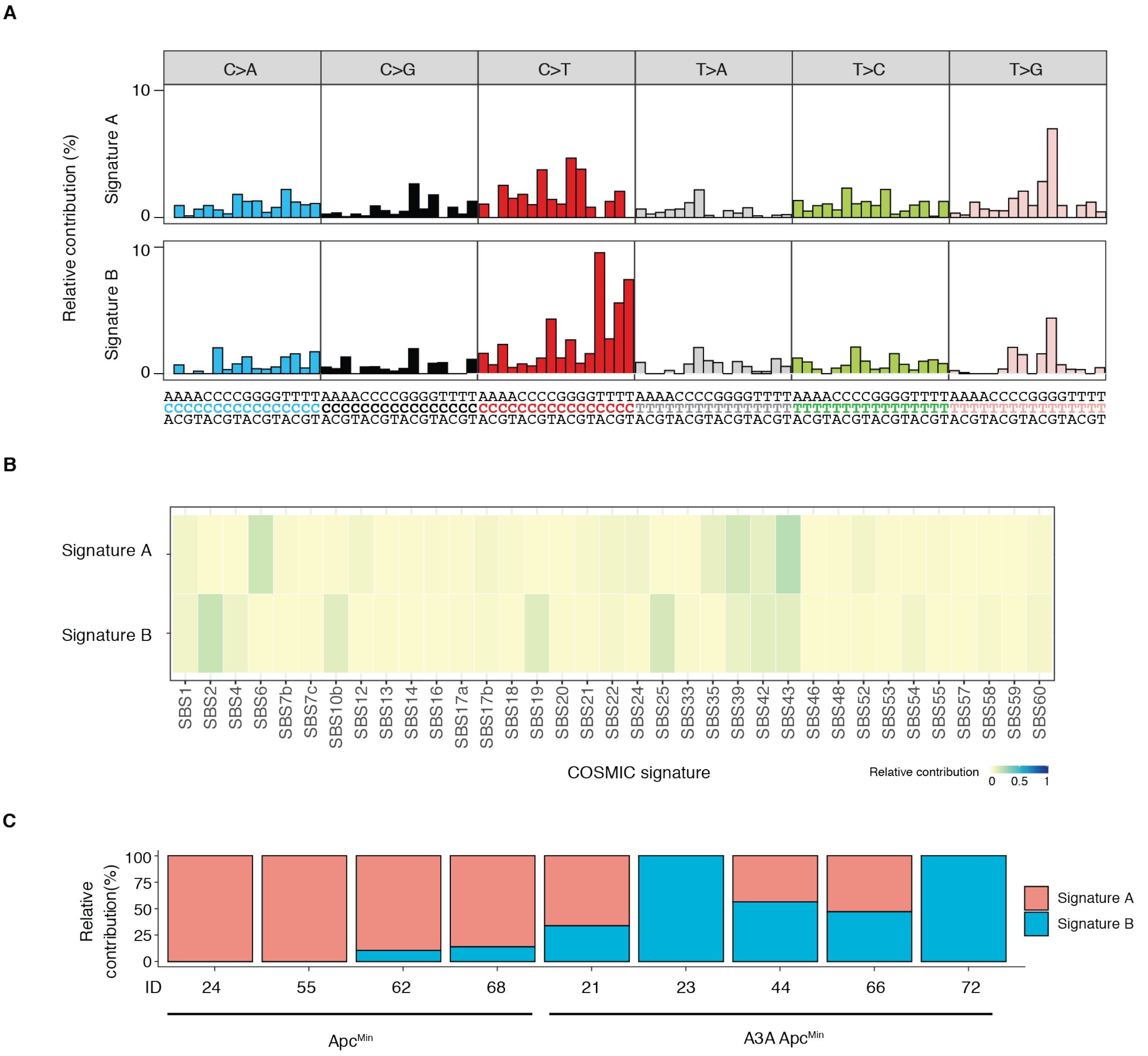


**Figure S8. *De-novo* non-negative matrix factorization (NMF) of predominant mutation signatures in exomes of mouse colorectal polyps.**

(**A**) Trinucleotide mutational frequency profiles of NMF-derived signatures arbitrarily designated "Signature A" and "Signature B".

(**B**) Heatmap denoting contribution of known COSMIC signatures to *de-novo* NMF-derived signatures. COSMIC signature SBS2 (APOBEC) is the most dominant feature of NMF-derived signature B. Only signatures with at least some detectable contribution are included.

(**C**) Stacked bar plots of the contribution of *de-novo* NMF signatures to mutational spectrum in Apc$^{Min}$ versus A3A$^{low}$/Apc$^{Min}$ tumors.

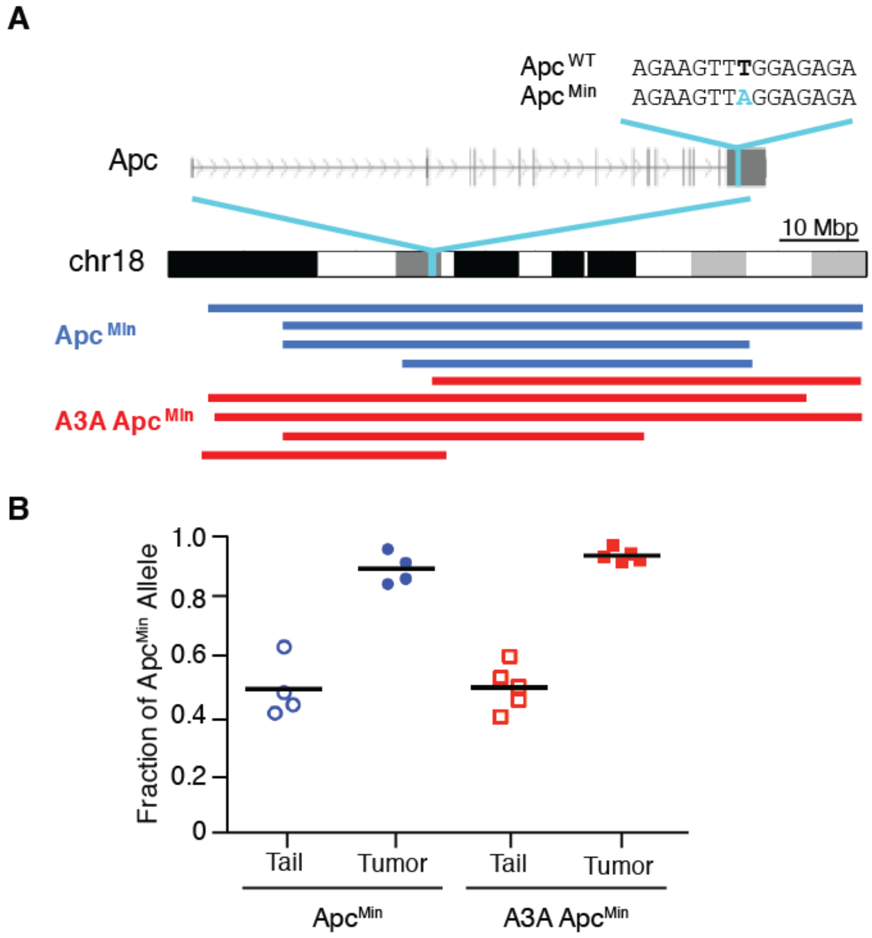


**Figure S9. Loss of heterozygosity (LOH) of Apc$^{Min}$ allele in colorectal tumors.**
(**A**) Ideogram of mouse chromosome 18. Locations of the *Apc* gene and the Apc$^{Min}$ mutation are marked in cyan. Horizontal blue (Apc$^{Min}$) and red (A3A/Apc$^{Min}$) lines represent chromosomal regions where LOH was detected in tumors in comparison to matched normal tissue.
(**B**) Dot plot showing the Apc$^{Min}$ allele frequency (chr18:34312601 T>A) in normal tail and tumor samples from Apc$^{Min}$ and A3A/Apc$^{Min}$ mice. Horizontal bar represents mean value.

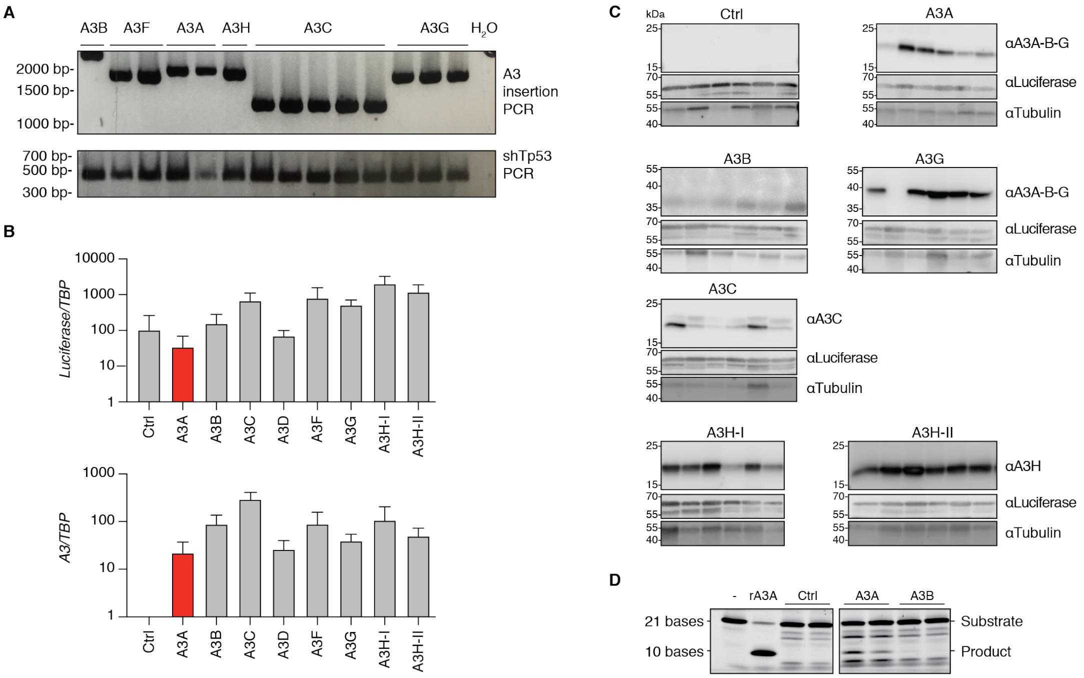


**Figure S10. APOBEC3 constructs are integrated and expressed in livers of injected *Fah*$^{-/-}$ mice.**

(**A**) PCR amplicons from genomic DNA extracted from livers of mice hydrodynamically injected with the indicated *A3* minigene constructs. This PCR assay uses primers flanking each differently sized *A3* minigene.

(**B**) RT-qPCR results for *Luciferase* mRNA (identical primers for all inserts) or specific *A3* mRNAs relative to those of the housekeeping gene *Tbp* in livers from hydrodynamically injected animals (mean +/- SD across all mice injected with the indicated *A3* minigene). Ctrl – control construct is identical apart from lacking an *A3* minigene.

(**C**) Representative immunoblots of liver protein extracts from animals of mice hydrodynamically injected with the indicated *A3* minigene constructs.

(**D**) Single-stranded DNA cytosine deamination activity of liver protein extracts from animals hydrodynamically injected with the indicated *A3* minigene constructs. Deamination substrate with no extract or with 1 nM purified recombinant (r)A3A are controls. A3A shows clear activity, and A3B shows weaker but still detectable activity.

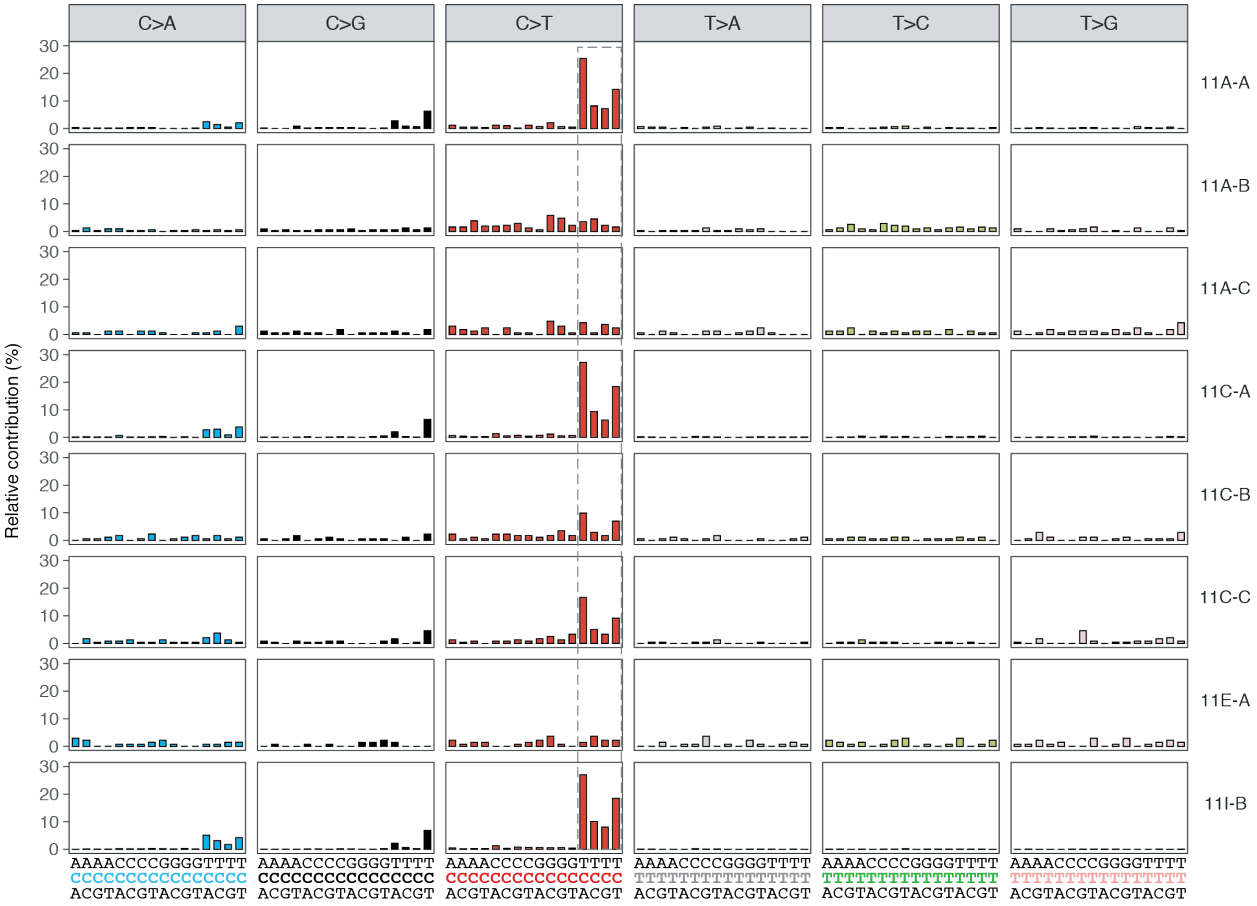


**Figure S11. Individual trinucleotide mutational profiles in liver tumors from A3A Fah mice.** Barplots representing composite trinucleotide mutation profiles for all base substitutions in exome-seq data from individual A3A Fah liver tumors. Dotted box highlights the trinucleotide motifs that define the APOBEC mutation signature.

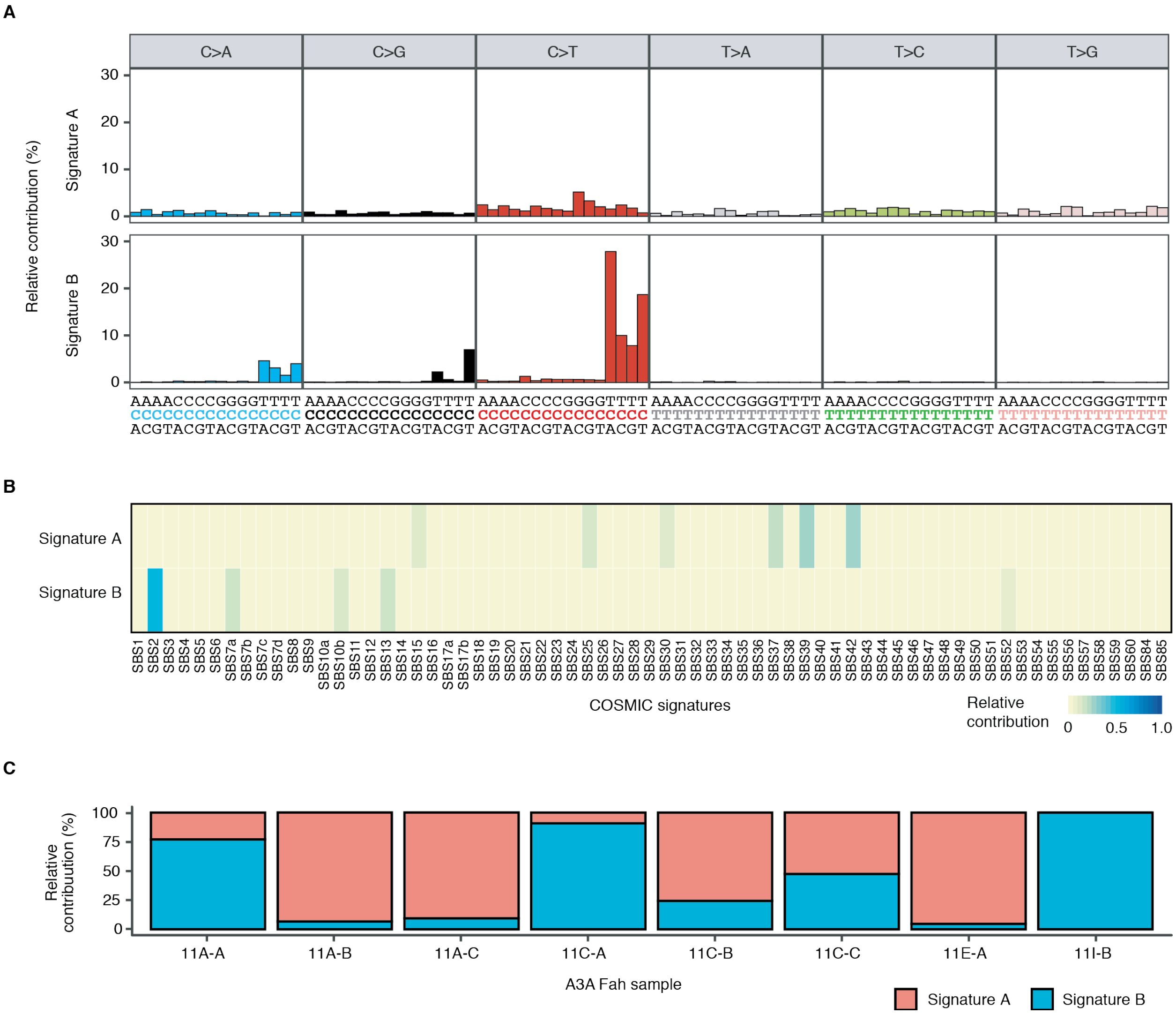


**Figure S12. *De-novo* non-negative matrix factorization (NMF) of predominant mutation signatures in exomes of liver tumors from A3A Fah mice.**

(**A**) Trinucleotide mutational frequency profiles of NMF-derived signatures arbitrarily designated "Signature A" and "Signature B".

(**B**) Heatmap denoting contribution of known COSMIC signatures to *de-novo* NMF-derived signatures. COSMIC signature SBS2 (APOBEC) is the most dominant feature of NMF-derived signature B.

(**C**) Stacked bar plots of the contribution of *de-novo* NMF signatures to mutational spectrum in individual A3A Fah tumors.
